# FINDER converts zero-background kinetic fingerprinting into area-scalable attomolar biomarker detection

**DOI:** 10.64898/2026.06.01.729299

**Authors:** Liuhan Dai, Pavel Banerjee, Alexander Johnson-Buck, Aaron Blanchard, Zi Li, Nils G. Walter

## Abstract

Background constrains analytical sensitivity: surveying larger sensor areas samples more analyte molecules but also accumulates false positives, limiting gains in detection performance. Here we introduce FINDER—Fluorogenic INstantaneous Digital Enumeration and Recognition—a single-molecule platform that combines kinetic fingerprinting with fluorogenic transient probes for rapid molecular classification under near-zero-background conditions. By suppressing both solution and surface-associated background at micromolar probe concentrations, FINDER classifies individual molecules within seconds-scale observation windows per field of view. This regime allows sensitivity to scale with surveyed sensor area, enabling amplification-free quantification of the miRNA cancer biomarker *hsa*-miR-16 with an 11 aM detection limit. FINDER further generalizes to HPV16 DNA biomarker detection, two-color RNA/DNA co-profiling, and rapid discrimination of clinically relevant EGFR single-nucleotide variants using multidimensional kinetic filtering. Rapid per-field classification permits tens of fields to be surveyed within minutes. By converting kinetic specificity into area-scalable sensitivity, FINDER enables semi-automated attomolar biomarker counting without amplification in practical workflows.

## Introduction

Digital biomarker assays based on imaging are widely used across modalities ranging from microarrays and immunofluorescence to single-molecule nucleic acid counting^1,2^. In these assays, target quantification ultimately reduces to distinguishing true, localized events from nonspecific background and then counting positives across the imaged sensor area^3,4^. Most platforms perform this discrimination by fluorescence intensity thresholds. However, because specific and nonspecific signals often occupy overlapping intensity distributions^5,6^, the threshold couples sensitivity and specificity: stringent thresholds suppress false positives but discard true events, whereas permissive thresholds improve detection recovery at the cost of increased background^7^. This limitation is particularly severe for amplification-free detection of nucleic acids, including circulating tumor DNA and microRNAs, where expected target counts are low and even rare false positives can dominate the measurement^8-13^.

Temporal information offers an orthogonal basis for molecular discrimination^14-17^. We and others have shown that kinetic fingerprinting can classify targets from time-resolved trajectories generated by transient probe interactions rather than by intensity alone, enabling rejection of off-target interactions and surface artifacts in a multidimensional kinetic feature space^12-19^. This capability is especially attractive for surface-based digital assays, because the number of true target detections increases linearly with surveyed area, or number *N* of fields of view (FOVs)^20-22^. In conventional non-zero background detection, however, false positives also accumulate linearly with area; so that the detection limit only improves as 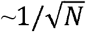. If false positives could be held near-zero while additional area if sampled, sensitivity would improve approximately linearly with *N*, providing a route to extreme sensitivity without enzymatic amplification^17,19^. Achieving this regime has remained elusive because accurate kinetic classification generally requires long observation windows for each FOV^17-19^, making large-area sampling impractically slow.

We therefore sought to accelerate kinetic fingerprinting while preserving near-zero background. High-confidence classification requires observing multiple (i.e., >10) binding and dissociation transitions at each molecular locus, so shorter acquisition times require acceleration of both association and dissociation. Unimolecular dissociation can be readily tuned through solution conditions, including temperature, ionic strength, and denaturants such as formamide^20^. Association, in contrast, is a bimolecular process and most directly accelerated by increasing the probe concentration^18,23-25^. Conventional fluorescent probes cannot be used at concentrations of >100 nM because unbound probes generate prohibitive background, even under total internal reflection fluorescence (TIRF) illumination^26-28^. Although zero-mode waveguides can enable single-molecule detection at elevated fluorophore concentrations, their fabrication typically requires specialized nanoscale lithography and serial manufacturing approaches that remain expensive and difficult to scale^29^. Fluorogenic probes that are quenched in solution and become fluorescent upon target binding^30,31^, provide a readily available means to increase probe concentration without increasing optical background^32-36^, but have not yet been considered for kinetic fingerprinting.

Here we present FINDER—Fluorogenic INstantaneous Digital Enumeration and Recognition—as a platform that combines transiently binding fluorogenic probes with kinetic fingerprinting to identify single molecules within seconds while maintaining near-zero background. By accelerating molecular classification of biomarkers, FINDER makes large-area kinetic sampling practical and converts kinetic specificity into scalable sensitivity. Using miRNA titrations, we validate linear areal scaling and achieve an amplification-free limit of detection of 11 aM. We further extend FINDER to viral DNA detection, chromatic co-profiling of RNA and DNA targets, and discrimination of clinically relevant single-nucleotide variants. Together, these capabilities establish FINDER as a generalizable route to attomolar, amplification-free digital diagnostics for liquid biopsy and related applications.

## Results

### Design principles for area-scalable kinetic fingerprinting

We designed FINDER to overcome the scaling limit imposed by residual false positives imaging-based detection assays. In conventional digital readouts, persistent fluorescent spots are called as positive or negative using a signal threshold^37,38^, so overlapping signal and background distributions often cause false positives that increase as additional FOVs are surveyed (Fig. 1a,b)^7^. FINDER instead identifies single molecular biomarkers from their temporal pattern of transient probe binding. Weak-affinity readout probes repeatedly associate with and dissociate from surface-tethered targets, producing fluorescence trajectories that can be classified using kinetic features such as transition number, dwell times and fluorescence off-times rather than intensity alone (Fig. 1a). This multidimensional classification provides a route to a zero-background detection (ZBD) regime where only true positive spots are counted (Fig. 1a, right)^17-19^.

**Fig. 1.**
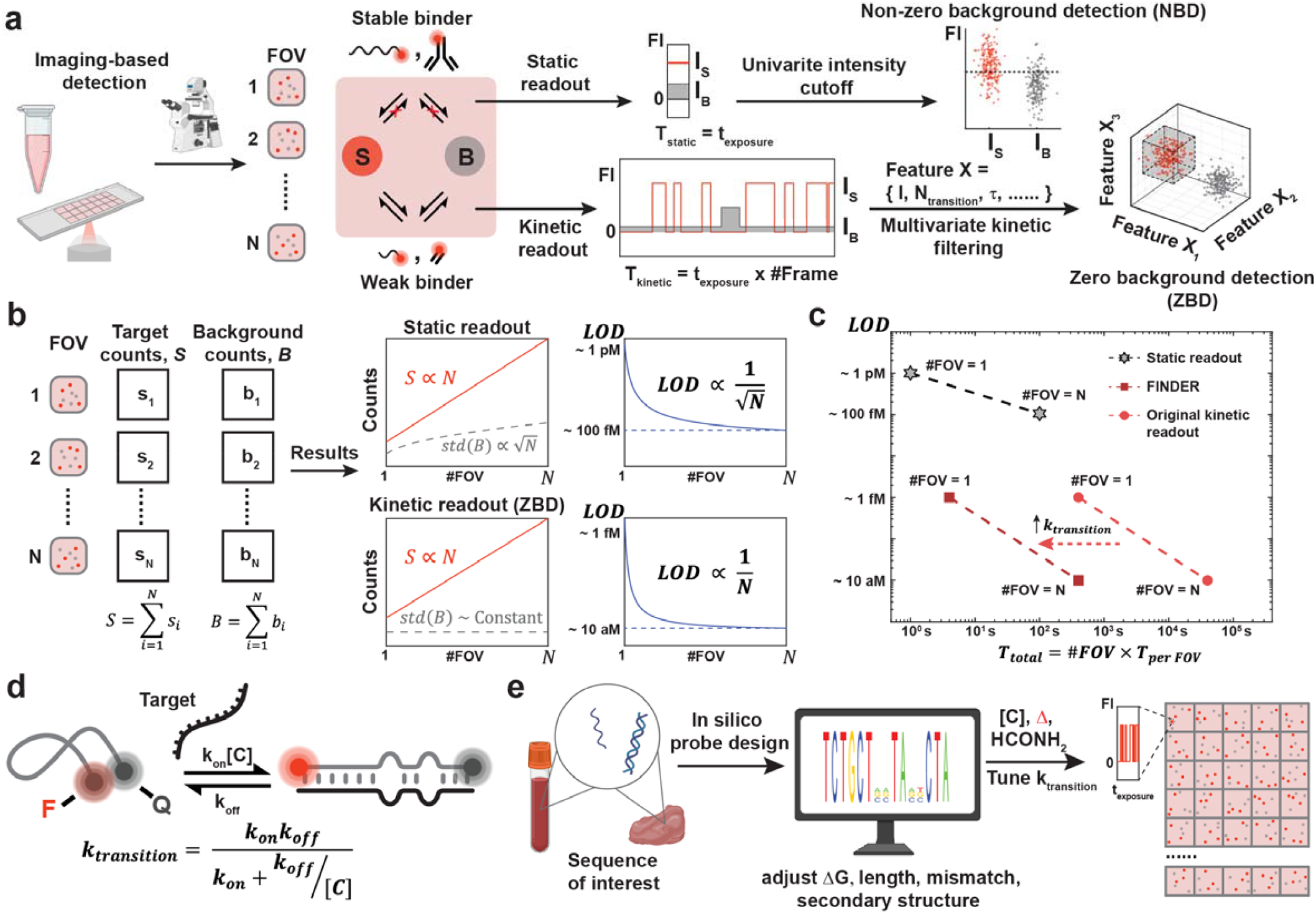
FINDER enables zero-background kinetic classification and areal scaling in digital imaging. **a**, Schematic comparing conventional static (intensity-thresholded) readout versus kinetic readout. Conventional readout utilizes static readout that relies on the intensity difference between target signals, *I*_*S*,_ and background signals, *I*_*B*_. In general, an optimal intensity cutoff is applied to separate *I*_*S*_ and *I*_*B*_ with a trade-off between false positives and false negatives. By contrast, kinetic readout uses a weaker binder that generates a molecular fingerprint of target molecules in time-series signals. Kinetic fingerprint is transformed into a multi-parameter kinetic space that allows complete separation between target signals (red dots) and off-target signals (gray dots) by applying a multi-parameter threshold (gray cube). This separation enables zero-background detection (ZBD). **b**, In imaging-based counting, accumulated counts across FOVs increase with the number of FOVs sampled. Under non-zero background detection (NBD; static thresholding), background counts accumulate with area and the limit of detection (LOD) improves approximately as 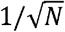. Under ZBD (FINDER), false-positive counts remain near-constant while true counts increase linearly with *N*, yielding an LOD that improves approximately as 1/*N*. **c**, Time–sensitivity trade-off for static readout, prior kinetic fingerprinting approaches, and FINDER. Increasing the kinetic transition rate (*k*_transition_) reduces the per-FOV observation time required to reach ZBD, making multi-FOV interrogation practical while preserving the areal-scaling benefit of ZBD. **d**, A dual-labeled probe (fluorophore–quencher) is quenched in solution and becomes fluorescent upon target binding, suppressing solution background and permitting micromolar imager concentrations to accelerate transitions. **e**, Candidate imagers are screened in silico for secondary structure and binding specificity and then experimentally optimized by tuning imager concentration, temperature and formamide to increase *k*_transition_while maintaining ZBD.

We formalized the sensitivity benefit of this regime using a Poisson model for digital counting across *N* FOVs or positions (Extended Data Fig. 1a). Under non-zero background detection (NBD), blank counts increase linearly with surveyed area, and the LOD decreases as 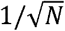 when defined by three standard deviations above the mean of the blank (Extended Data Fig. 1b,c)^9^. In contrast, in the ZBD limit, false-positive counts remain approximately constant as area increases, whereas true target counts continue to accumulate linearly. This allows the LOD to approach 1/*N* scaling with imaged area (Fig. 1b; Extended Data Fig. 1c). The practical challenge is throughput: previous kinetic fingerprinting approaches required long observation windows per FOV, preventing sampling of large sensor surface areas (Fig. 1c)^12,17,22^.

To make area-scalable kinetic fingerprinting practical, FINDER uses fluorogenic readout probes that support high probe concentrations while suppressing solution-phase fluorescence background (Fig. 1d). Each probe is dual-labeled with a fluorophore and a quencher, remaining dim in solution and becoming fluorescent upon target binding^26-28,32,34,36^. This binding-induced light-up design permits probe concentrations in the micromolar range, which accelerates biomarker association—i.e., increase the number of observable binding transitions within a short acquisition window—while maintaining sufficient contrast for single-molecule detection. Dissociation rates were tuned by engineering probe mismatches to permit transient binding despite forming 11 base pairs with the target across a high-specificity ∼13-nucleotide footprint, and the imaging conditions, including temperature and formamide concentration^24,25^, to generate trajectories suitable for rapid classification.

We paired this probe architecture with an in silico probe screening and optimization workflow for rapid adaptation to new nucleic acid targets following common thermodynamic principles^24,25^ (Fig. 1e; Methods; Supplementary Note 1). Candidate probes were prioritized for low secondary structure, favorable binding kinetics and specificity margins against related sequences, followed by empirical tuning of probe concentration and solution conditions to maximize transition rates while maintaining near-zero background. Using this framework, we developed FINDER probes for miRNAs, viral DNA and oncogenic single-nucleotide variants, establishing a general design framework for scalable kinetic digital detection (Supplementary Table 1).

### Linear areal scaling enables rapid low-attomolar miRNA detection

MicroRNAs are established circulating biomarkers, but their low abundance, short length, and vulnerability to assay background create a need for amplification-free methods capable of detecting very low copy number^10,11^. Human microRNA-16, or *hsa*-miR-16, stands out as one of the best-studied mammalian microRNAs with broad clinical relevance in cancer and cardiovascular disease because it regulates cell-cycle progression, apoptosis, inflammation, and stress responses^39,40^. We used *hsa*-miR-16 as a test case to experimentally validate the central premise of FINDER: when background is maintained near-zero, sensitivity should improve with imaged area. Using our probe workflow, we designed a fluorogenic imager for *hsa*-miR-16 (Fig. 2a; Extended Data Fig. 2a). Comparison with a conventional, non-fluorogenic version showed that, fluorogenicity was essential for operation at micromolar probe concentrations: at 1 µM, conventional fluorescent imagers resulted in prohibitive solution background, whereas fluorogenic imagers preserved single-molecule contrast and accelerated switching kinetics, thereby increasing the number of transitions per locus (Supplementary Fig. 1)^26-28,34,36^. However, blank controls with micromolar fluorogenic imagers also revealed numerous long-lived, bright surface-associated spots (Extended Data Fig. 2b; Supplementary Fig. 2). Their long-step-like intensity trajectories were consistent with nonspecific surface trapping and subsequent photobleaching rather than transient target binding (Supplementary Fig. 2).

**Fig. 2.**
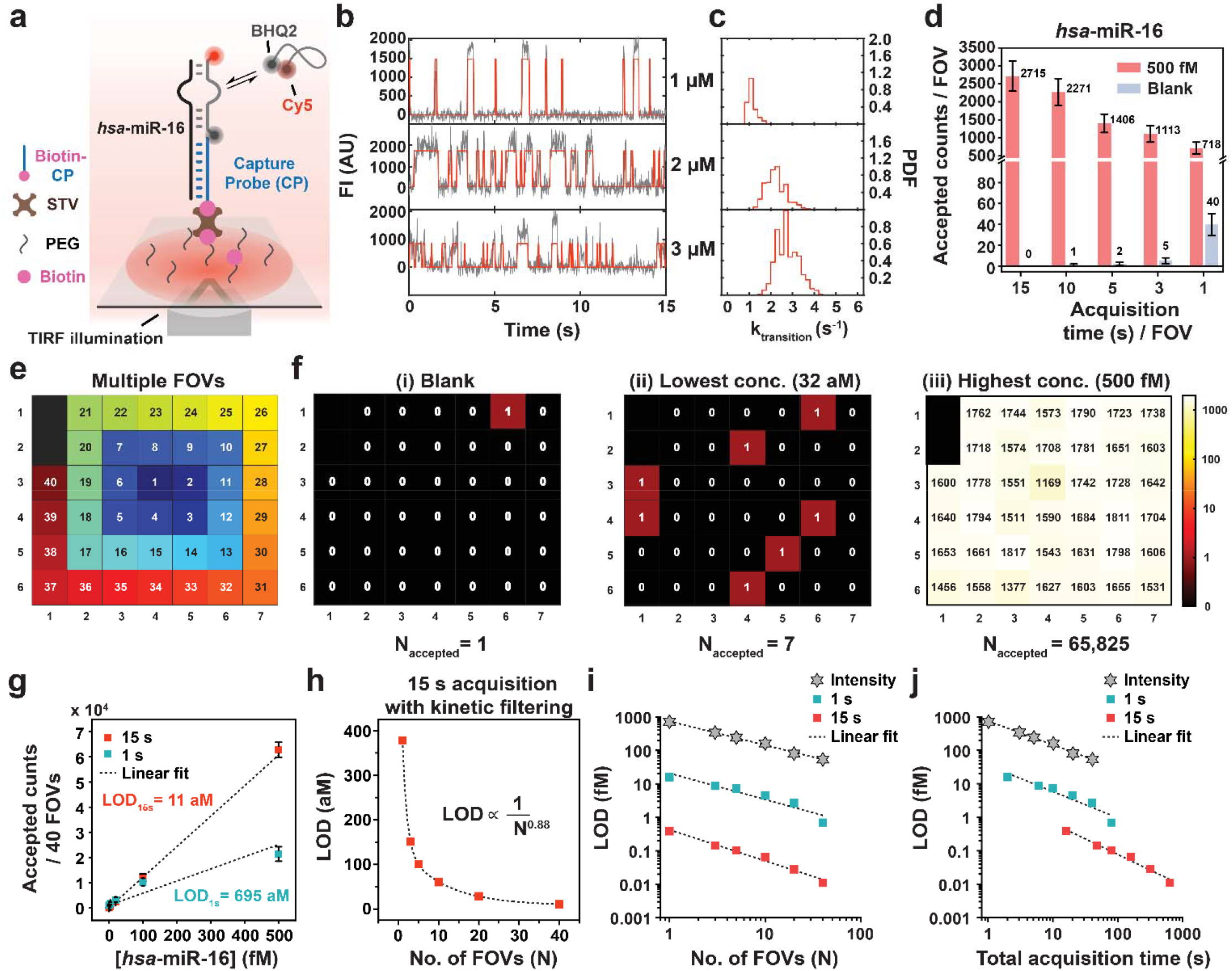
FINDER enables linear areal scaling and attomolar miRNA detection with seconds-per-field kinetic readout. **a**, FINDER assay schematic for *hsa*-miR-16 where a PEG-passivated, streptavidin-functionalized surface captures targets via hybridization to a biotinylated capture probe (CP). Objective-type TIRF excites surface-associated fluorogenic imagers to record kinetic fingerprints. **b**, Representative single-molecule trajectories at the indicated fluorogenic imager concentrations. **c**, Corresponding *k*_transition_distributions for the conditions in **b. d**, Accepted counts per FOV for 500 fM *hsa*-miR-16 and blank at the indicated acquisition times (2 µM imager). Black error bars indicate standard errors of the mean from three independent replicates. **e**, Spiral stage trajectory used to acquire 40 FOVs. **f**, Spatial maps of accepted counts across 40 FOVs for Blank, lowest concentration (32 aM) and highest concentration (500 fM). **g**, Calibration curves from 40 FOVs comparing 15-s and 1-s acquisition per FOV. **h**, LOD as a function of the number of FOVs imaged for 15-s acquisition, demonstrating near-inverse scaling with area. **i**, Comparison of LOD scaling across intensity thresholding, 15-s FINDER and 1-s FINDER as a function of FOV number. **j**, Comparison of LOD scaling versus total acquisition time for the detection schemes in **i**.

To suppress these false positives associated with fluorogenic probes, we introduced a quencher probe blocker (QPB): a short (5 nt), quencher-labeled oligonucleotide designed to bind and quench the open form of the fluorogenic imager while not impending its binding to the target miRNA (Extended Data Fig. 2a). As predicted, in blank samples, QPB reduced long-lived events and shifted the distribution of maximum bound times toward shorter durations (Extended Data Fig. 2b,c). In target-containing samples, it also improved the signal-to-noise ratio (SNR) of accepted traces by further reducing the ensemble fluorescence from unbound imagers (Extended Data Fig. 2d,e). We next optimized the imager concentration to balance transition rate against SNR. Increasing imager concentration increased the observed transition rate *k*_transition_ (Fig. 2b,c; Extended Data Fig. 2f) but also decreased SNR (Extended Data Fig. 2g). Using an empirical trace-acceptance criterion of SNR ≥ 3, we selected 2 µM as the optimal concentration for subsequent experiments (Extended Data Fig. 2g).

We next determined the per-FOV observation time required to achieve near-zero false positives under these conditions. Guided by the relationship between transition number, observation time and target-identification confidence (Supplementary Note 2), we tested a series of acquisition windows at 2 µM imager. Shorter acquisition times increased blank acceptance, consistent with trajectories being too short to reliably distinguish target from blank kinetic signatures. In contrast, 15 s per FOV reduced blank events to near-zero while preserving high target counts (Fig. 2d). This observation window is an order of magnitude shorter than earlier monochromatic implementations of kinetic fingerprinting, which typically required minutes per FOV to establish a definitive kinetic signature^12,17,22^, and is sufficiently brief to enable practical multi-FOV sampling. Using 15 s per FOV, we imaged 40 FOVs in a spiral acquisition pattern (Fig. 2e). Accepted target counts increased proportionally with the number of FOVs surveyed, whereas blank counts remained nearly constant (Fig. 2f). In an miRNA titration series, this areal sampling strategy enabled amplification-free detection of *hsa*-miR-16 at an LOD of 11 aM (Fig. 2g). Consistent with the areal-scaling model, the LOD improved approximately inversely with the number of FOVs imaged in the 15-s regime (Fig. 2h).

In contrast, shortening the acquisition to 1 s per FOV increased blank acceptance and degraded classification, yielding a substantially higher LOD of 695 aM and scaling behavior approaching the 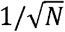 non-zero-background regime (Fig. 2g,i; Extended Data Fig. 2h–l). Intensity thresholding showed a similarly sublinear scaling with FOV number and a much higher LOD in the femtomolar range (Fig. 2i; Extended Data Fig. 2j–l), consistent with the accumulation of that blank counts when events are called using intensity alone^9^. Together, these results define the practical conditions required for areal scaling in FINDER. Kinetic classification improves sensitivity two orders of magnitude relative to intensity thresholding, even in fast-acquisition regimes; however, attomolar performance requires per-FOV observation time to maintain near-zero background, allowing multi-FOV sampling to translate directly into lower LODs (Fig. 2i,j).

### Generalizing FINDER to viral DNA detection and two-color kinetic co-profiling

To establish that FINDER extends beyond miRNA detection and can support parallel multi-analyte measurements, we next targeted human papillomavirus type 16 genomic DNA (HPV16), a clinically important biomarker associated with cervical cancer and the increasing incidence of HPV16-positive head-and-neck cancers^41^. We designed a fluorogenic imager and an LNA-modified capture probe to stabilize surface tethering under elevated temperature and denaturant conditions used to tune hybridization kinetics (Fig. 3a; Extended Data Fig. 3a). Initial HPV16 measurements produced fewer accepted counts than expected and showed poor separation from blanks, indicating inefficient target capture (Extended Data Fig. 3b,c). Assay performance depended strongly on capture conditions likely reflecting reduced target accessibility caused by secondary structure or self-interactions within the capture configuration. Increasing temperature and/or adding formamide improved the kinetic separation of HPV16 and blank measurements (Extended Data Fig. 3b,c). We therefore selected 37 °C without formamide, which maintained high accepted counts with low blank acceptance (Extended Data Fig. 3b,c), for subsequent HPV16 experiments.

**Fig. 3.**
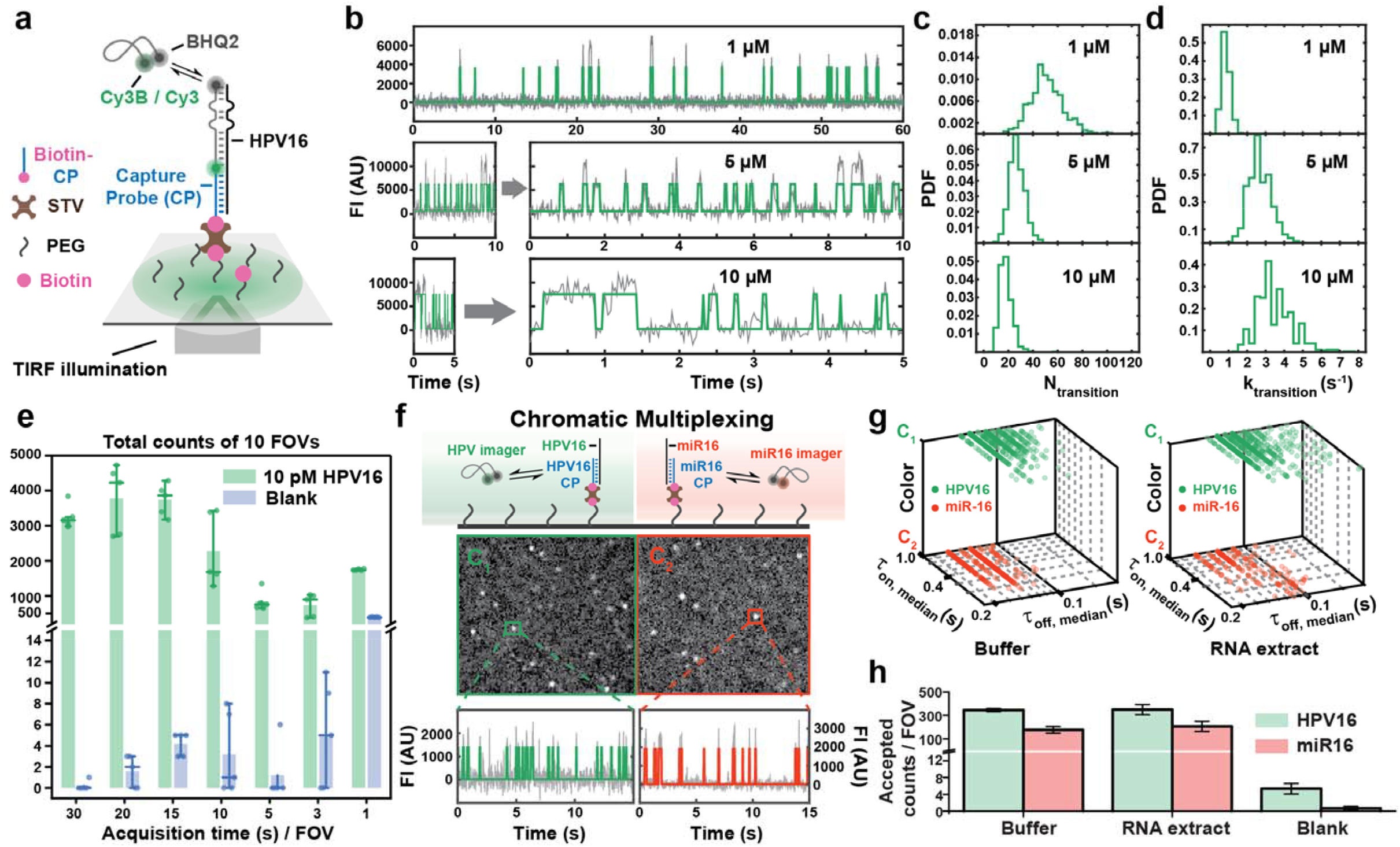
FINDER detects HPV16 and enables chromatic multiplexing of HPV16 and *hsa*-miR-16. **a**, FINDER assay schematic for HPV16 detection using a surface-tethered capture probe and a fluorogenic imager. **b**, Representative single-molecule fluorescence trajectories at the indicated imager concentrations. FI, fluorescence intensity. AU, arbitrary unit. **c**,**d**, Distributions of the number of transitions (*N*_transition_; **c**) and transition rate (*k*_transition_; **d**) for the conditions in **b. e**, Total accepted counts across 10-FOV detection of HPV16 at 5 µM imagers with different acquisition time per FOV. Each datapoint is total counts across 10 FOVs generated by applying a set of kinetic filtering criteria using custom-built optimizer program. Column height means total counts. n = 5 rounds of kinetic filtering criteria optimizations. Middle line is median total count. Whiskers represent the IQR from 5% to 95%. **f**, Two-color (chromatic) kinetic multiplexing scheme for simultaneous detection of HPV16 and *hsa*-miR-16 (top) and representative images (middle) and example trajectories (bottom) from the HPV16 (green) and *hsa*-miR-16 (red) channels are shown. **g**, Separation of HPV16 and *hsa*-miR-16 populations in a multidimensional kinetic feature space (color channel, *τ*_on,median_and *τ*_off,median_) in buffer and RNA extract. **h**, Accepted counts per FOV for HPV16 and *hsa*-miR-16 in buffer or, RNA extract, and a no-target blank in buffer. Black error bars indicate standard errors of the mean from three independent replicates.

We next optimized imaging conditions to accelerate HPV16 target sampling while maintaining trace quality. Increasing the fluorogenic imager concentration from 1 to 10 µM increased transition statistics and *k*_transition_ (Fig. 3b–d; Extended Data Fig. 3d), but higher concentrations reduced SNR for a substantial fraction of candidate trajectories (Extended Data Fig. 3e). This behavior is consistent with the concentration–SNR trade-off intrinsic to single-molecule fluorescence measurements^26-28^. We therefore selected 5 µM as an operating concentration that enhanced transition rates while maintaining preserving acceptable trace SNR (Fig. 3b–d). We then evaluated acquisition time requirements across 10 FOVs. Extending the observation window progressively reduced blank acceptance without compromising HPV16 counts, and 30 s per FOV yielded near-zero blank acceptance across the 10-FOV dataset (Fig. 3e). Although longer than the acquisition time required for *hsa*-miR-16, this per-FOV observation window remains compatible with multi-FOV sampling and supports practical areal scaling for viral DNA detection.

Finally, we used FINDER to demonstrate chromatic kinetic co-profiling^13^, configuring a two-color assay for simultaneous detection of HPV16 DNA and *hsa*-miR-16. Both capture probes were combined on the same surface and interrogated with spectrally distinct imagers (Fig. 3f). Time-series acquisition resolved the two targets as separable populations in a joint feature space defined by channel color and dwell-time statistics (Fig. 3g). Using the same analysis pipeline, we detected both targets in buffer and in RNA extract background with minimal cross-channel interference (Fig. 3h). The HPV16 channel showed a modest increase in blank acceptance in the two-color format relative to the HPV16 single-plex condition (Fig. 3h versus Fig. 3e), consistent with additional nonspecific events when both imagers and capture probes are present in the same well. Even so, background remained low relative to the on-target signal and did not preclude detection of either target under the conditions tested.

### Multidimensional kinetics enable rapid discrimination of clinically relevant SNVs

Single-nucleotide variants (SNVs) present a stringent specificity challenge for amplification-free nucleic acid detection: mutant and wild-type sequences differ by only a single base, while clinically relevant mutant alleles can be outnumbered by wild-type background. Sensitive and quantitative detection of oncogenic *EGFR* mutations in circulating tumor DNA is therefore a key target application for digital assays^9,12,18,42^. We tested whether FINDER could achieve second-scale SNVs discrimination by coupling fluorogenic readout to multidimensional kinetic classification. As an initial test case, we targeted the *EGFR* T790M mutation and designed a fluorogenic imager to maximize thermodynamic and kinetic contrast between mutant T790M and wild-type T790 sequences (Fig. 4a; Extended Data Fig. 4a). To increase the number of informative transition within short observation windows, we tuned hybridization kinetics using formamide, temperature and imager concentration^23-25^. Adding 10% formamide shortened bound times and increased *k* _transition_, estimated for each trajectory by exponential fitting of HMM-derived dwell-time statistics (Fig. 4b,c; Extended Data Fig. 4b; Methods). These observations are consistent with formamide destabilizing probe-target duplexes and accelerating exchange at fixed probe concentration^24^. Because higher probe concentrations tend to further increase transition rates but reduce single-molecule SNR, we compared fluorophore-quencher pairs to maximize fluorogenic contrast. Cy3B–BHQ2 outperformed Cy5–BHQ2 in both single-molecule SNR and ensemble fluorogenic ratio measurements (Fig. 4d,e; Supplementary Fig. 3) and was used for subsequent experiments.

**Fig. 4.**
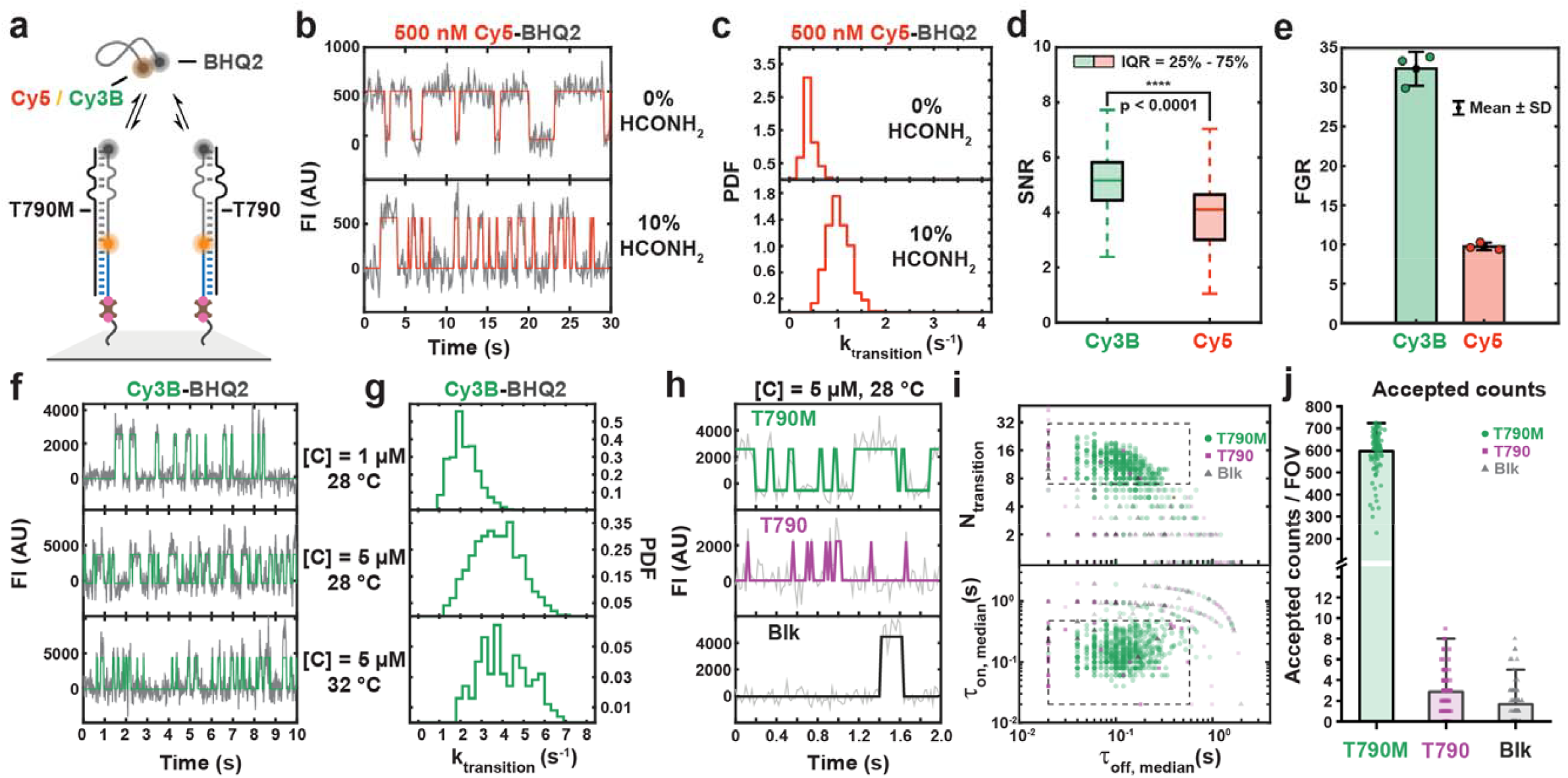
FINDER enables ultrafast discrimination of SNV T790M in *EGFR* Exon 20. **a**, Assay schematic for distinguishing *EGFR* T790M from wild-type T790 using a fluorogenic imager. **b**, Representative single-molecule trajectories acquired with 0% or 10% formamide. **c**, *k*_transition_distributions for the conditions in **b. d**, Signal-to-noise ratio (SNR) distributions for Cy3B- and Cy5-labeled imagers. Boxes indicate the interquartile range; center lines indicate the median; whiskers indicate the 5th–95th percentiles. P-value is shown in the panel (statistical test described in Methods). **e**, Ensemble fluorogenic ratio (FGR) for Cy3B and Cy5 imagers (mean ± s.d.; n=3 independent experiments). **f**, Representative trajectories at the indicated imaging conditions used to increase transition rates while maintaining classification performance. **g**, *k*_transition_distributions for the conditions in **f. h**, Representative trajectories for T790M, T790 and blank acquired at the selected condition with 2-s observation per FOV. **i**, Separation of T790M, T790 and blank in a multidimensional kinetic feature space defined by *N*_transition_, *τ*_on,median_and *τ*_off,median_. Dashed lines indicate kinetic filtering thresholds. **j**, Accepted counts per FOV for T790M, T790 and blank with 2-s observation (*n* = 86 FOVs). Boxes indicate the interquartile range; center lines indicate the median; whiskers indicate 5th–95th percentiles.

We next optimized imaging conditions to maximize transition sampling while maintaining trace quality. Increasing the imager concentration from 1 to 5 µM increased transition rates (Fig. 4f,g). In contrast, elevating temperature over the tested range shortened dwell times but did not increase the net transition rate, because unbound intervals lengthened concurrently, with some binding events now too short at our time resolution (Extended Data Fig. 4c). We therefore selected 5 µM imager at 28 °C in 10% formamide for rapid classification (Fig. 4f,g). Under these conditions, FINDER distinguished T790M from wild-type T790 with only 2 s of observation per FOV. Representative trajectories revealed distinct switching behavior for mutant, wild-type and blank samples (Fig. 4h). In a multidimensional kinetic feature space defined by *N*_transition_, *τ*_on,median_and *τ*_off,median_, predefined thresholds separated T790M biomarkers from wild-type and blank distributions (Fig. 4i). Across 86 FOVs, accepted counts were significantly higher for T790M than for wild-type and blank controls (Fig. 4j), demonstrating that multidimensional kinetic filtering preserves single-base specificity even during only seconds-per-FOV acquisition.

To ask whether this strategy generalizes to other SNV, we designed and optimized an assay for a second clinically relevant *EGFR* mutation, L858R^18^. Using an LNA-modified capture probe and the same in silico workflow, we identified conditions that produced distinct kinetic fingerprints for L858R, wild-type L858 and blank samples (Extended Data Fig. 4d–f). Increasing formamide and temperature accelerated exchange while reducing residual wild-type interactions to near-blank levels under the selected imaging conditions (Extended Data Fig. 4e-h). These optimized conditions separated L858R from wild-type and blank controls within a 10-s observation window (Extended Data Fig. 4i,j; Supplementary Fig. 4). Together, these results establish that FINDER can be tuned for rapid, amplification-free discrimination of clinically important SNVs through fluorogenic probes and multidimensional kinetic classification.

## Discussion

FINDER establishes a practical route to ultrahigh-sensitivity, amplification-free digital imaging by coupling multidimensional kinetic classification for false-positive suppression with fluorogenic probe engineering that minimizes unbound-probe fluorescence and shortens the time needed for classification sufficiently to achieve areal scaling. In this near-zero-background regime, surveying additional fields of view increases analyte counts without a corresponding accumulation of blank events, allowing sensitivity gains to approach 1/*N* areal scaling rather than the 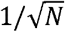 scaling imposed by non-zero background. Using this principle, FINDER achieves an amplification-free limit of detection of 11 aM for *hsa*-miR-16 and generalizes to viral DNA detection, chromatic RNA–DNA co-profiling, and rapid SNV discrimination with 2-30 s acquisition times per FOV. Together, these results establish kinetic fingerprinting as a broadly applicable strategy for high-specificity, single-molecule counting across biomarker classes^12,13,17-19,42^.

FINDER’s central enabling advance is the fluorogenic readout architecture, which suppresses solution background and thus permits micromolar probe concentrations that increase transition statistics without sacrificing single-molecule contrast^26-28^. This speed is more than a throughput improvement; it makes multi-FOV kinetic classification practical and therefore operationalizes areal scaling. In parallel, the probe design and optimization workflow provides a reproducible path from sequence selection to assay conditions across distinct nucleic-acid targets, including miRNA, viral DNA and mutant DNA, supporting the platform’s generality. Notably, in contrast to another recently published approach to generating fluorogenic probes for super-resolution imaging^36^, the engineered mismatches of FINDER probes permit transient binding despite forming 11 base pairs with the target over an ∼13-nucleotide footprint, thereby increasing target specificity relative to conventional 8-9 nucleotide SiMREPS probes^17-22^.

FINDER is readily amenable to further engineering. First, performance depends on surface passivation quality; at micromolar probe concentrations, imperfect PEGylation can result in trapped or adsorbed probe populations that mimic true target binding events. We mitigated this effect for miRNA detection using a quencher probe blocker, QPB, and extending QPB-style suppression to additional targets should further improve blank performance and enable even more robust areal scaling. Second, the per-field observation time required to maintain near-zero background is target- and condition-dependent; for HPV, longer acquisition windows were required than for *hsa*-miR-16, likely reflecting differences in capture efficiency, probe kinetics, and matrix effects. Third, the current workflow still relies on deliberate parameterization of kinetic filters and empirical tuning of temperature, formamide and probe concentration. More automated approaches—including robust classifier training on labeled kinetic trajectories and kinetically informed computational sequence design—could further reduce user burden, improve portability across instruments and sample types^43,44^, and enable multiplexing without forcing compromises among the optimal conditions for different targets.

Looking ahead, FINDER provides a foundation for assays with higher multiplexing and broader analyte scope. We demonstrated two-color co-detection of RNA and DNA targets; combining chromatic multiplexing with iterative probe exchange could expand panels to dozens of biomarkers while preserving the benefits of amplification-free quantification^5,7,13^. Beyond nucleic acids, extending FINDER to proteins will require programmable weak binders with fluorogenic behavior. Emerging strategies for rapidly engineered protein and peptide binders may provide a route to develop such reagents, potentially enabling a unified, amplification-free digital platform for multiomic diagnostics^15,45^. More broadly, by transforming kinetic specificity into sensitivity linearly scaling with sensor area, FINDER maximizes the analytical advantage of large-area imaging and establishes a practical path toward attomolar digital diagnostics for liquid biopsy and related applications.

## Methods

### Materials

DNA and RNA oligonucleotides were obtained from Integrated DNA Technologies (IDT) with HPLC purification. Sequences and modifications are provided in Supplementary Table 1. Unless stated otherwise, oligonucleotides were reconstituted in TE (10 mM Tris-HCl, 0.1 mM EDTA, pH 8.0) to 100 µM, aliquoted to avoid repeated freeze–thaw cycles, and stored at −20 °C (DNA) or −80 °C (RNA). Reagents for buffer preparation were purchased from Sigma-Aldrich or Thermo Fisher Scientific. DNA LoBind tubes (Eppendorf) were used throughout to reduce adsorption losses.

### Preparation of glass slide surfaces for single-molecule microscopy

Glass coverslips (No. 1.5, 24 × 50 mm; VWR, 48393-241) were passivated with polyethylene glycol (PEG) using NHS-ester chemistry. Briefly, a 1:100 molar mixture of biotin–PEG–SVA and mPEG–SVA (Laysan Bio; Biotin-PEG-SVA-5K-100MG and MPEG-SVA-5000-1g) was applied following established cleaning and aminosilanization (using (3-aminopropyl) triethoxysilane, Sigma-Aldrich, 440140-100ML) procedures^12,13^. After modification, coverslips were stored protected from light (aluminum foil) in a nitrogen-purged cabinet and used within 4 weeks. For imaging, multiwell sample chambers were assembled by trimming P20 micropipette tips (Thermo Fisher Scientific, 02-682-261) to retain the wide end as a well. The cut tips were placed wide-end down onto the PEGylated coverslip and sealed with epoxy (Ellsworth Adhesives, 4001).

### Design of FINDER readout probes (fluorogenic imagers) and capture probes

Fluorogenic readout probes (“imagers”) were designed using an in silico screening workflow that generates candidate sequences complementary to the target site, prioritizes low predicted secondary structure and self-dimerization, and scores candidates based on binding thermodynamics and predicted specificity against plausible off-target sequences. Candidate imagers were further evaluated by secondary-structure prediction under assay-relevant ionic strength and temperature conditions (NUPACK). Design details, scoring criteria and parameter choices are provided in Supplementary Note 1.

Capture probes were designed to stably tether targets under the most stringent capture/imaging conditions used in this study. Where required, locked nucleic acid (LNA) modifications were incorporated to maintain target binding in the presence of elevated temperature and formamide; formamide effects on duplex stability were considered during probe selection and condition optimization^24^.

### FINDER assay workflow (target capture and kinetic fingerprinting)

Unless stated otherwise, all steps were performed at room temperature. PEG-passivated sample wells were prepared as described above. Wells were washed once with T50 buffer (10 mM Tris-HCl, pH 8.0, 50 mM NaCl) and incubated with streptavidin (40 µl, 1 mg ml□^1^ in T50) for 10 min. Wells were then washed three times with 1× PBS. Biotinylated capture probes (100 nM in 1× PBS) were thermally conditioned by heating to 85 °C for 5 min, incubating at 37 °C for 5 min and cooling to room temperature to disrupt any self-association or secondary structure, then added to wells for 10 min to functionalize the surface. After three washes with 4× PBS, targets were introduced at the indicated concentrations in 4× PBS supplemented with 2 µM poly(dT) carrier oligonucleotide. For *hsa*-miR-16, target capture was performed for 1 h at room temperature without formamide. For other targets, capture was performed for 1 h at the indicated temperature with or without formamide, as specified for each assay (Supplementary Table 2). Wells were covered to minimize evaporation during incubations. After capture, wells were washed three times with 4× PBS.

Immediately before imaging, 50–100 µl of imaging buffer containing fluorogenic imagers at the indicated concentration was added together with an oxygen scavenging system (OSS): 1 mM Trolox, 5 mM protocatechuic acid (PCA) and 50 nM protocatechuate 3,4-dioxygenase (PCD). Samples were then imaged by objective-type TIRF microscopy. Stock solutions were prepared as follows: PCD stock was prepared in 100 mM Tris-HCl, pH 8.0, 50 mM KCl, 1 mM EDTA and 50% (v/v) glycerol; PCA was prepared as a 100 mM solution in water and adjusted to pH 8.3 with KOH; Trolox was dissolved in water and adjusted to pH 10–11 with KOH. Stock solutions were aliquoted and stored at −20 °C.

### Single-molecule fluorescence microscopy

Most single-molecule measurements were performed on an Oxford Nanoimager (ONI) configured for objective-type total internal reflection fluorescence (TIRF) microscopy (instrument specifications available from the manufacturer). Imaging was conducted using a 100× 1.4 NA oil-immersion objective with the built-in Z-lock autofocus enabled. Because the ONI’s internal temperature regulation is limited near ambient conditions, prolonged laser illumination can lead to gradual warming and thermal drift. To stabilize temperature during acquisition, the instrument housing was externally cooled by circulating water from a temperature-controlled bath through a metal heat-sink assembly clamped to the microscope body. The bath setpoint was maintained 5–6 °C below the target sample temperature to stabilize imaging temperature during acquisition. Unless stated otherwise, *hsa*-miR-16 experiments were acquired using 640 nm excitation (Cy5) at 20 ms exposure per frame. HPV16 experiments were acquired using 532 nm excitation (Cy3/Cy3B) at 20 ms exposure per frame. For two-color acquisitions, 532 nm and 640 nm were used for Cy3/Cy3B and Cy5, respectively, with simultaneous dual-channel acquisition. For multi-field acquisition, automated stage scanning was controlled using a custom Python script via the ONI application programming interface.

To maximize imaged area for *hsa*-miR-16 standard curves and LOD/scaling measurements (Fig. 2g–j), data were acquired on an Olympus IX83 microscope equipped with a 100× 1.5 NA oil-immersion objective (Olympus UPlanApo), an Abbelight SAFe360 fiber-coupled illuminator and a Hamamatsu ORCA-Fusion 4.0 sCMOS camera. Temperature was maintained at 25 °C using an objective heater (Bioptechs) and a stage-top incubator (Tokai Hit). Samples were excited at 647 nm; imaging was performed over a 1280 × 1280 pixel region of interest corresponding to ∼124 µm × 124 µm in the sample plane at 20 ms exposure per frame. Automated stage scanning was used to acquire 40 fields of view per sample well in a spiral pattern (Fig. 2e). Data processing and kinetic filtering were performed as described below and in Supplementary Table 3.

### Multiwell sample chips for large-field measurements

For large-field measurements, multiwell sample chips were fabricated by laminating 6.35-mm-thick black acrylic (McMaster-Carr, 8505K754) onto double-sided adhesive (3M, 8153LE), laser cutting 80 mm × 24 mm chips with 18 wells (3.8 mm diameter) and bonding them to 50 mm × 24 mm biotin-PEG/mPEG-coated coverslips. Reagent additions followed the same workflow as for standard wells, using reduced volumes to match the well geometry (20 µL for streptavidin and capture probe solutions; 50 µL for washes and imaging solutions).

### Analysis of single-molecule data

Single-molecule movies were analyzed using custom MATLAB scripts following established diffraction-limited kinetic fingerprinting workflows^17,18^. Briefly, candidate binding sites were identified within each FOV by computing an intensity-fluctuation map (mean absolute frame-to-frame intensity change per pixel across the movie) and locating local maxima in the fluctuation map. For each maximum, a 3 × 3-pixel region of interest (ROI) was defined and converted into a background-subtracted integrated intensity trajectory.

Trajectories were idealized using a two-state hidden Markov model (HMM) implemented in a customized vbFRET-based pipeline^17,18^. From HMM-idealized traces, kinetic features were extracted, including *N*_transition_, *τ*_on,median_, *τ*_off,median_and maximum dwell-time metrics where indicated. Signal-to-noise ratio (SNR) was computed as the standard deviation of the trajectory divided by the mean intensity difference between bound and unbound states. Events were classified as accepted by applying thresholds on intensity, SNR and kinetic features; the general parameter set is provided in Supplementary Table 3.

### Chromatic multiplexing and kinetic feature-space visualization

For two-color experiments, capture probes for HPV16 and *hsa*-miR-16 were incubated together on the same PEG–streptavidin surface (total capture probe concentration 100 nM in PBS). Targets (or target mixtures) were introduced in the appropriate capture buffer and incubated for 1 h. For complex matrix experiments, RNA extract was diluted 100-fold in capture buffer before incubation^13^. After capture, sample wells were washed three times with PBS and exchanged into imaging buffer containing both fluorogenic imagers simultaneously (Cy3/Cy3B channel for HPV16 and Cy5 channel for *hsa*-miR-16) at the concentrations used for the corresponding single-plex assays. Time-series movies were then acquired by objective-type TIRF microscopy with sequential or split-channel detection, as configured on the instrument.

For visualization and classification of multiplexed data, kinetic features extracted from each accepted trajectory (for example, *τ*_on,median_and *τ*_off,median_) were plotted in a two-parameter kinetic space with color channel encoded as an additional dimension, enabling separation of HPV16 and *hsa*-miR-16 populations in a 3D scatter representation (Fig. 3g).

### Fitting of the cumulative frequency of dwell times

Dwell times were extracted from HMM-idealized trajectories and analyzed in MATLAB. Bound- and unbound-state dwell times were pooled separately for each condition and binned at the camera integration time (20 ms). The complementary cumulative distribution of bound or unbound dwell times was fit with a single-exponential model,

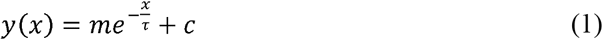

where *x* is dwell time and *m, τ* and *c* are fit parameters. For normalized curves spanning 1 to 0, *m*was fixed to −1; *c* captures any long-time offset and *τ* is the characteristic (mean) dwell time. Fits were retained when the sum of squared errors was < 0.05 and *R*^2^>0.99.

### Limit of detection (LOD) estimation

Calibration curves were generated by imaging serial dilutions of synthetic targets and quantifying accepted event counts after kinetic filtering. For *hsa*-miR-16 (Fig. 2g), counts were summed across 40 fields of view (FOVs) per chamber for each concentration (15-s and 1-s acquisition per FOV as indicated). A linear regression was fit to accepted counts versus concentration over the range used for quantification. The limit of detection (LOD) was then calculated using a blank-based criterion:

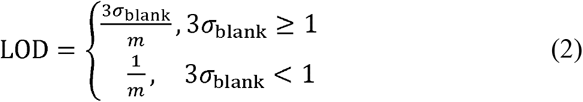

where *σ*_blank_ is the standard deviation of total accepted counts across 40 FOVs measured in blank controls processed and analyzed identically to samples, and *m*is the slope of the calibration curve (accepted counts per concentration, using the same 40-FOV aggregation).

For LOD-versus-FOV scaling analyses (Fig. 2h–j and Extended Data Fig. 2i,k), LOD values for a given number of FOVs (*N*) were estimated by random subsampling without replacement from the full multi-FOV dataset using custom MATLAB code. For each *N*, we randomly selected *N*FOVs, summed accepted counts across the selected FOVs for blank and sample conditions, and computed the corresponding LOD using the equation above (with *σ*_blank_ and *M* evaluated for that same *N*). This subsampling procedure was repeated at least 10 times for each *N*; the plotted LOD values reflect the mean across resamples (and variability across resamples where indicated).

### Statistical tests

Boxplots show the interquartile range (IQR) with the median as the center line and whiskers spanning the 5th–95th percentiles, as indicated in figure legends. Statistical significance was evaluated using two-sided Student’s t-tests (unpaired unless stated otherwise), and the corresponding P-values are reported in the figures. Experimental conditions (temperature, pH and salt/formamide concentration) were controlled as described in Methods.

## Data availability

The data supporting the findings of this study are available within this Article and its Supplementary Information. Source data are provided with this paper. Any additional data are available from the corresponding author upon request.

## Code availability

The code used in this study has been described in reference^13,17^ and is available through a DeepBlue repository. Any additional information concerning the code is available from the corresponding author upon request.

## Acknowledgements

We thank Sujay Ray (University of Mississippi) for helpful discussions and Muneesh Tewari (University of Michigan) for guidance in selecting HPV16 as a target. This work was supported by the NIH grants R21 CA225493 and R35 GM131922 (N.G.W.).

## Author contributions

L.D., P.B. and N.G.W. conceived the study. L.D. and P.B. designed and performed most experiments. A.B. and Z.L. performed initial quencher-screening experiments informing the fluorogenic probe design. A.B. developed the fluorogenic probe design code. L.D., P.B. and A.J.B. analyzed the data and prepared the figures. L.D. and P.B. wrote the initial manuscript draft. N.G.W. was responsible for the conception and supervision of the project. All authors reviewed and edited the manuscript.

## Additional information

### Supplementary information

Supplementary information is available in the online version of the paper.

**Correspondence and requests for materials** should be addressed to Nils G. Walter.

---

**Extended Data Fig. 1.**
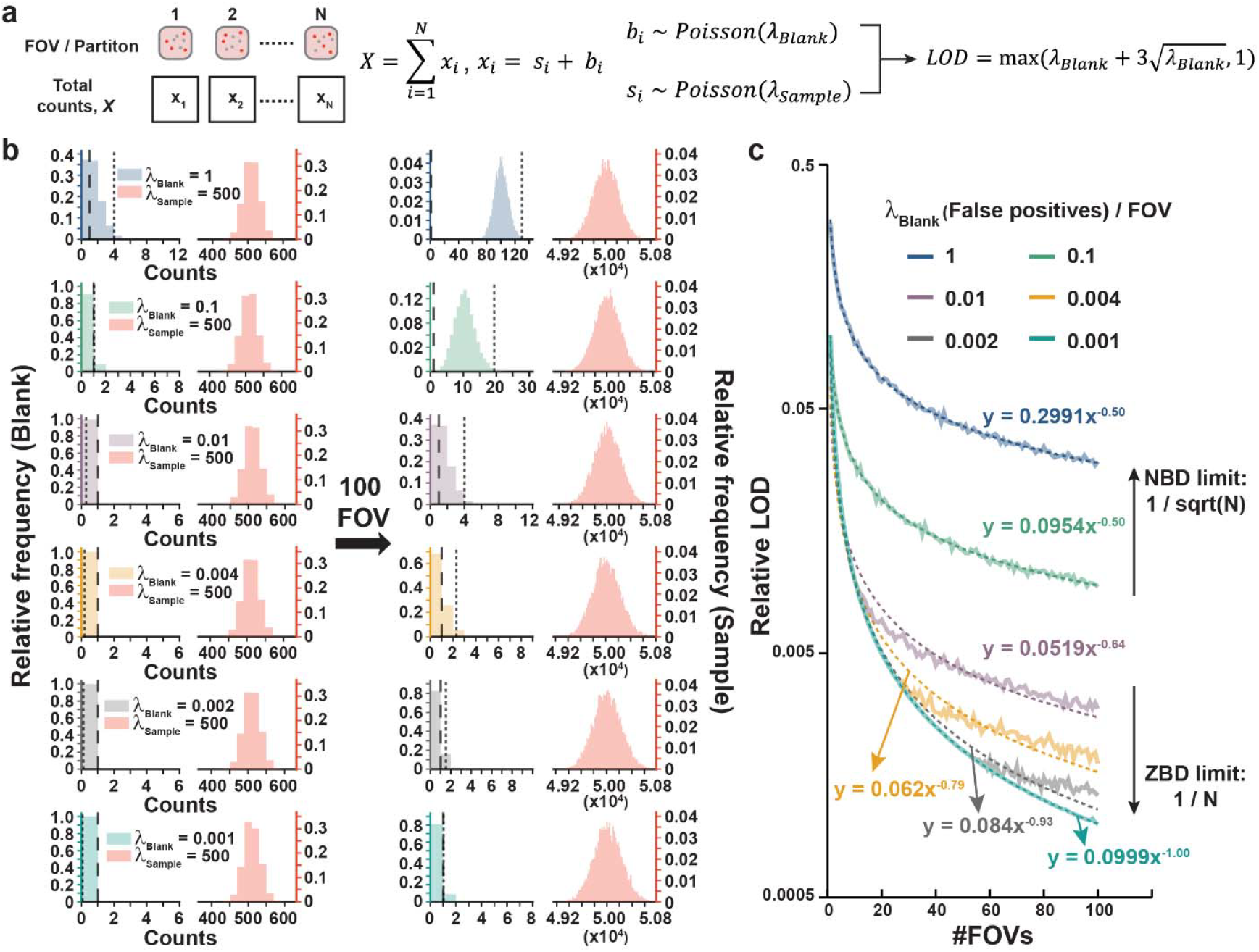
Poisson simulation of areal scaling under non-zero and zero-background detection. **a**, Poisson counting model for accumulated signal and background across N FOVs or partitions. Total counts are 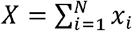, where *x*_*i*_ = *s*_*i*_ + *b*_*i*_, *b*_*i*_ = Poisson (*λ*_blank)_and *s*_*i*_ = Poisson(*λ*_sample_). The detection threshold is defined as 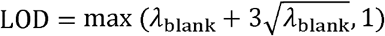, which accounts for cases where the 3-standard-deviation threshold is less than one positive count per measurement. **b**, Example Poisson count distributions for blank and sample at different *λ*_blank_ values for a single FOV (left) and for *N* = 100 FOVs (right). Counts are binned into unit-width intervals (*k, k* + 1), such that an integer count *k* contributes to the bin from *k* to *k*+ 1. Dashed lines indicate the threshold set by 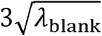 (small separation) or by 1 count (large separation). **c**, Simulated relative LOD versus number of FOVs for different false-positive rates per FOV (*λ*_blank_). Solid curves show simulation results and dashed curves show power-law fits. When background counts accumulate (non-zero background detection, NBD), LOD improves approximately as 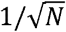; when false positives are suppressed (zero-background detection, ZBD), LOD approaches 1/*N* scaling.

**Extended Data Fig. 2.**
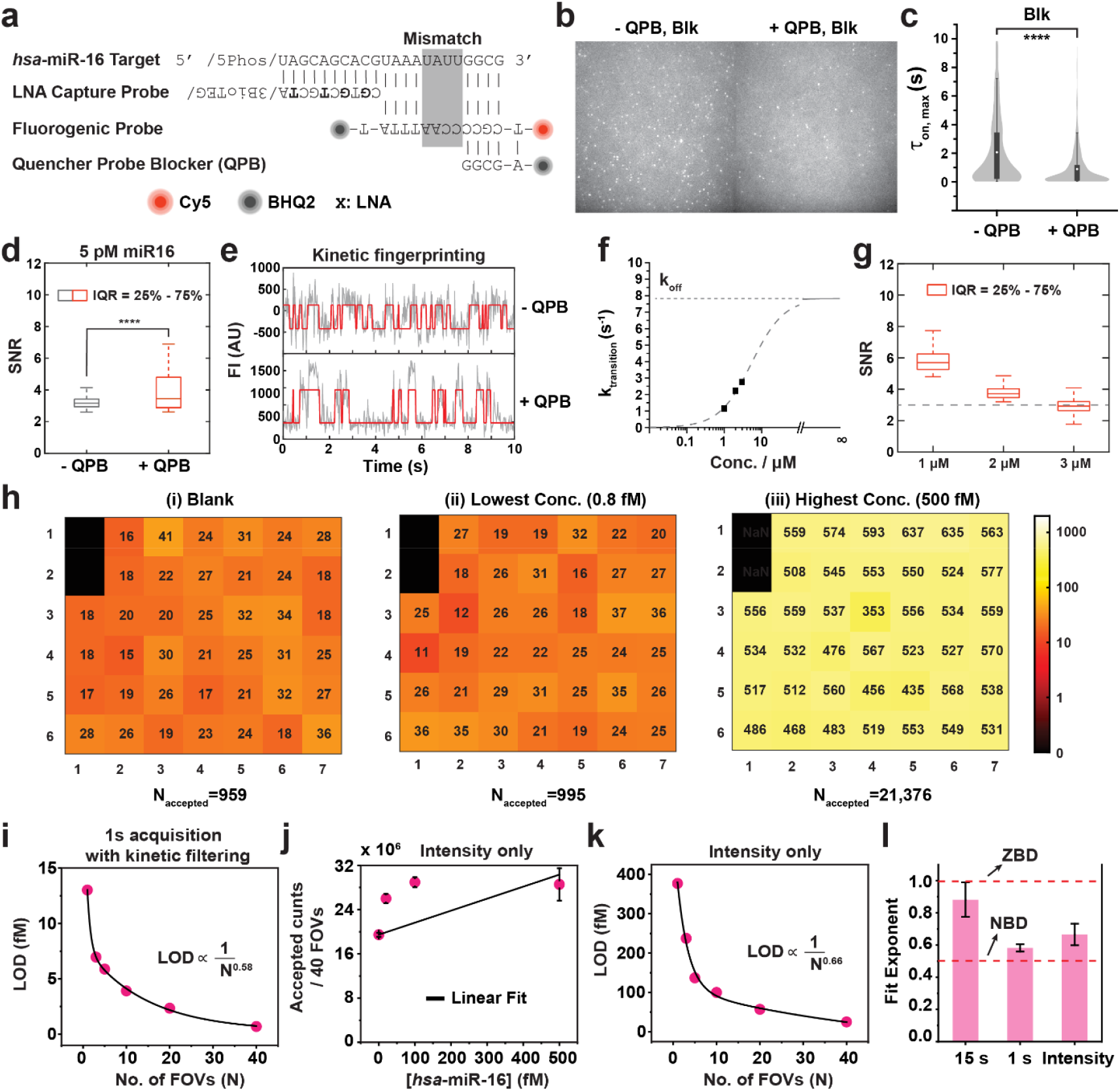
Optimization and scaling behavior for *hsa*-miR-16 detection using FINDER. **a**, Oligonucleotide design for *hsa*-miR-16 detection, including the LNA-modified capture probe, fluorogenic imager and quencher probe blocker (QPB). Imager–target mismatches are indicated. **b**, Representative blank images acquired without and with QPB. **c**, Distribution of the maximum bound time (*τ*_on,max_) for candidate events in blanks without and with QPB. Boxes indicate the interquartile range; center lines indicate the median; whiskers indicate the 5th–95^th^ percentiles, ****P < 0.001. **d**, SNR distributions for accepted traces at 5 pM *hsa*-miR-16 without and with QPB. Boxplot definitions as in **c. e**, Representative accepted kinetic trajectories corresponding to **d** (gray, raw; red, HMM idealization). **f**, Estimated transition rate (*k*_transition_) as a function of imager concentration with extrapolation to the dissociation-rate limit (*k*_off_). **g**, SNR distributions for traces passing intensity thresholding at the indicated imager concentrations; dashed line indicates SNR = 3. **h**, Example 40-field acquisition with 1-s observation per field. Heat maps show accepted counts after kinetic filtering for (i) blank, (ii) 0.8 fM and (iii) 500 fM concentration. **i**, LOD versus number of FOVs for 1-s acquisition with kinetic filtering; solid line shows a power-law fit. **j**, Accepted counts versus *hsa*-miR-16 concentration using intensity thresholding (40 FOVs); line shows a linear fit. **k**, LOD versus number of FOVs for intensity thresholding; solid line shows a power-law fit. **l**, Fit exponents for LOD scaling under 15-s FINDER, 1-s FINDER and intensity thresholding. Error bars indicate s.d. across independent experiments (*n*=3 experiments).

**Extended Data Fig. 3.**
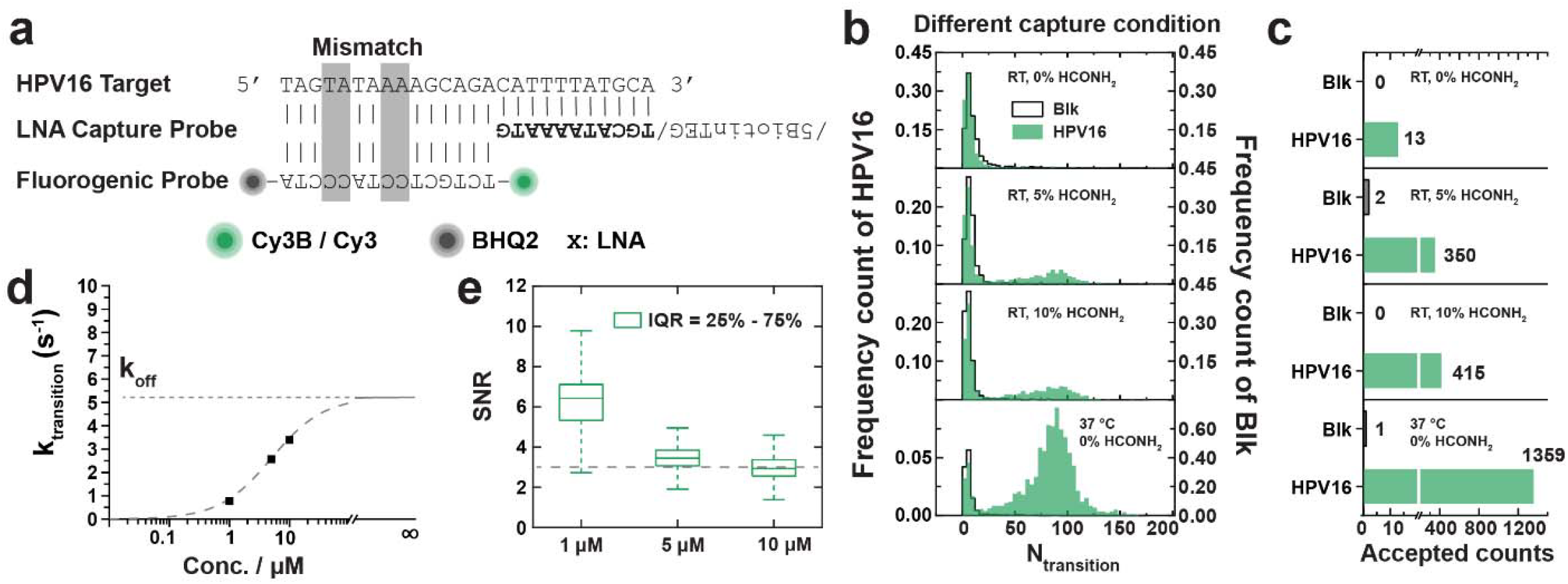
Optimization of HPV16 detection using FINDER. **a**, Oligonucleotide design for HPV16 detection, including the LNA-modified capture probe and fluorogenic imager. Imager–target mismatches are indicated. **b**, Distributions of transition counts (*N*_transition_) for blank and 10 pM HPV16 under the indicated capture conditions. RT, room temperature. **c**, Accepted counts for blank and HPV16 under the capture conditions in **b** after applying kinetic filtering. **d**, Estimated transition rate (*k*_transition_) as a function of imager concentration with extrapolation to the dissociation-rate limit (*k*_off_). **e**, SNR distributions for traces passing intensity thresholding at the indicated imager concentrations; dashed line indicates SNR = 3.

**Extended Data Fig. 4.**
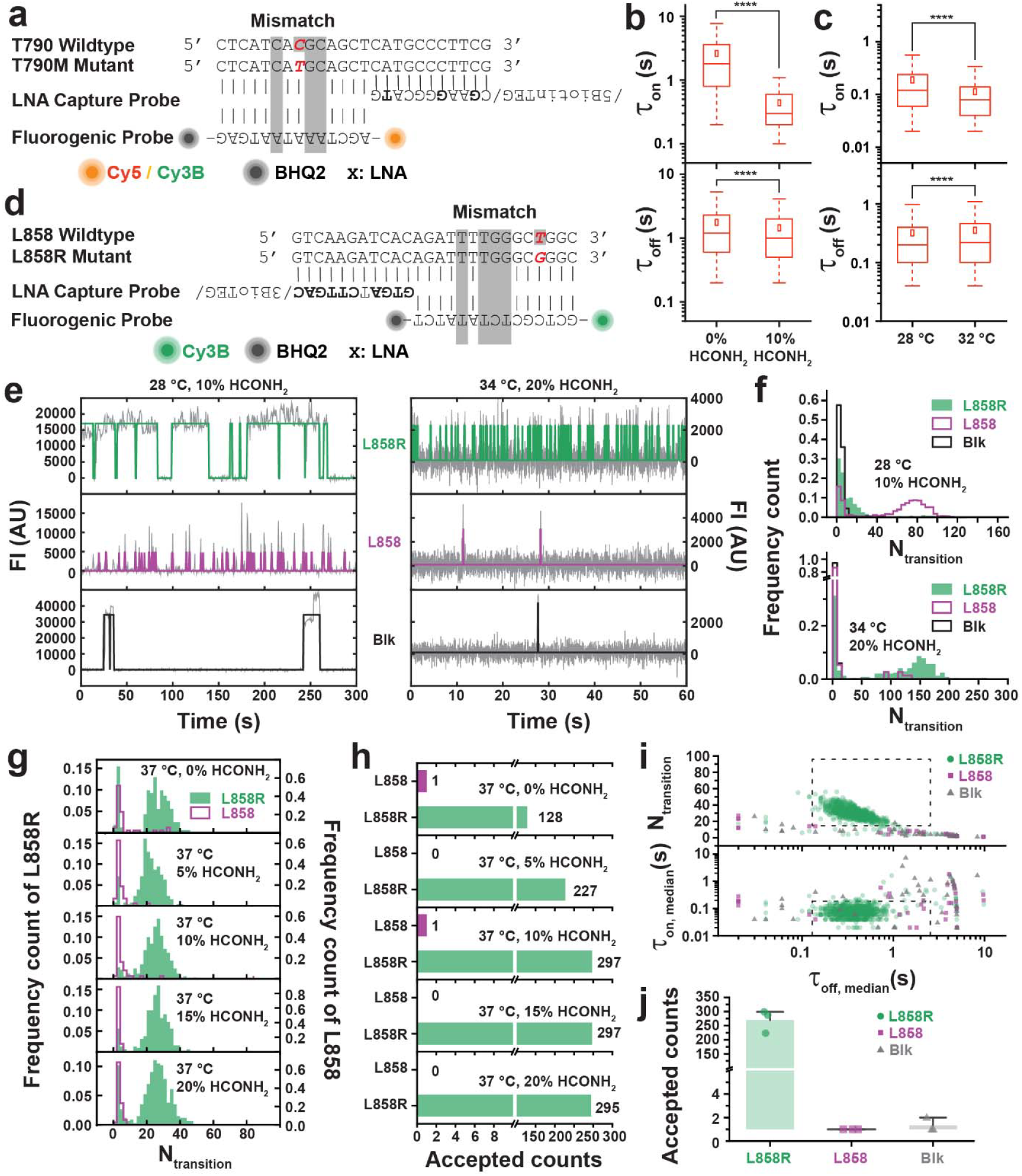
Assay design and condition optimization for SNV detection (*EGFR* T790M and L858R) using FINDER. **a**, Oligonucleotide design for *EGFR* T790M detection, including the LNA-modified capture probe and fluorogenic imager. Imager–target mismatches are indicated. **b**, Effects of formamide on dwell times (*τ*_on_ and *τ*_off_) for the T790M assay. Boxes indicate the interquartile range; center lines indicate the median; whiskers indicate the 5th–95th percentiles, ****P < 0.001. **c**, Effects of temperature on dwell times for the T790M assay. Boxplot definitions as in **b. d**, Oligonucleotide design for *EGFR* L858R detection, including the LNA-modified capture probe and fluorogenic imager. Imager–target mismatches are indicated. **e**, Representative single-molecule trajectories for L858R (mutant), L858 (wild type) and blank under the indicated imaging conditions. **f**, *N*_transition_distributions for blank, 10 pM L858 and 10 pM L858R under the imaging conditions in **e. g**, *N*_transition_distributions for 10 pM L858 and 10 pM L858R under the indicated capture conditions. **h**, Accepted counts for L858 and L858R after applying kinetic filtering for the capture conditions in **g. i**, Separation of L858R, L858 and blank in a multidimensional kinetic feature space defined by *N*_transition_, *τ*_on,median_, and *τ*_off,median_. Dashed lines indicate kinetic filtering thresholds. **j**, Accepted counts per FOV for L858R, L858 and blank with 10-s observation. Points denote individual FOVs; bars denote mean. Error bars indicate 1 s.d.

