## Supplementary Figures, Tables and Notes for "FINDER converts zero-background kinetic fingerprinting into area-scalable attomolar biomarker detection"

#### TABLE OF CONTENTS

Table of Contents ……………………………………………………………….…………1

Supplementary Figures……………………….………………………………………......4

Supplementary Fig. 1 | Fluorogenic imagers suppress solution background and accelerate kinetic readout for *hsa*-miR-16 …………………………………………………………………...4

Supplementary Fig. 2 | Representative traces from non-specifically surface-trapped fluorogenic imagers in the absence of quencher probe blocker (QPB)……………………………………...6

Supplementary Fig. 3 | Ensemble fluorogenicity of imagers labeled with different fluorophores…7

Supplementary Fig. 4 | Representative kinetic trajectories for *EGFR* L858R, wild-type L858 and blank………………………………………………………………………………………….8

Supplementary Note 1 | Computational pipeline for fluorogenic probe design for FINDER…….13

Part 1: High-Throughput Screening of Candidate Sequences (*HTfluorogenicScreenV2_3.m*)…..13

1.1 Initialization and Parameter Definition…………………………………………………...13

1.2 Target Sequence Definition…………………………………………………………….14

1.3 Reference Thermodynamic Calculation…………………………………………………..14

3.3 Iterative Optimization……………………………………………………………………..20

Supplementary Note 2 | Determination of required imaging time from a target significance level.22

Supplementary References………………………………………………………………..24

#### Supplementary Figures

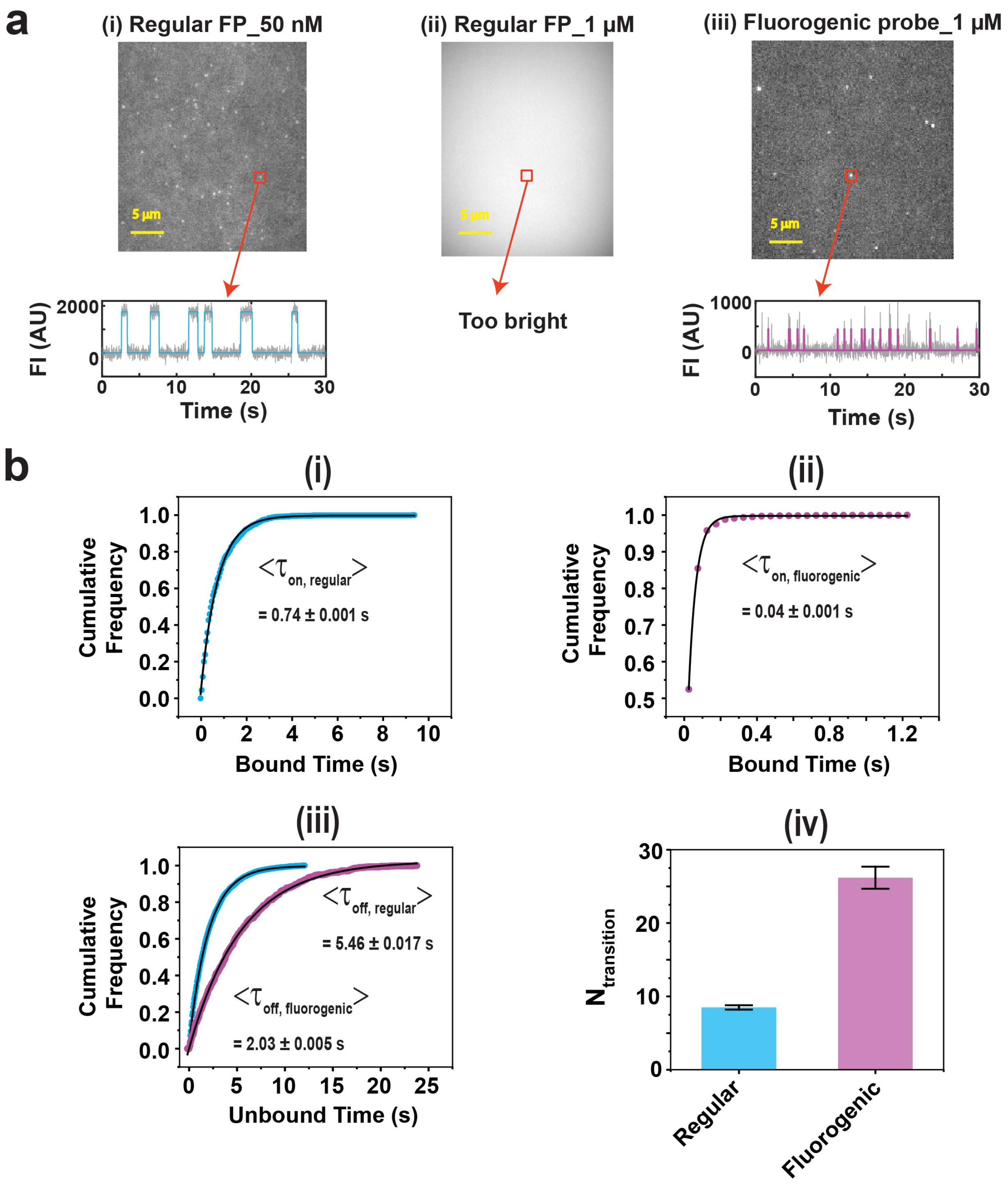

**Supplementary Fig. 1 | Fluorogenic imagers suppress solution background and accelerate kinetic readout for *hsa*-miR-16. a,** Representative images and single-molecule trajectories comparing regular fluorescent imagers and fluorogenic imagers. With regular imagers, increasing probe concentration from 50 nM to 1 μM produces prohibitive solution background, whereas fluorogenic imagers permit micromolar probe concentrations while maintaining single-molecule contrast. **b,** Dwell-time and transition statistics comparing regular and fluorogenic imagers. **(i,ii)** Cumulative distributions of bound times ($\tau_{\text{on}}$) with exponential fits. **(iii)** Cumulative distributions of unbound times ($\tau_{\text{off}}$) with exponential fits. **(iv)** Mean number of transitions ($N_{\text{transition}}$) per accepted trace. Error bars indicate s.d. across independent experiments ($n = 3$ experiments).

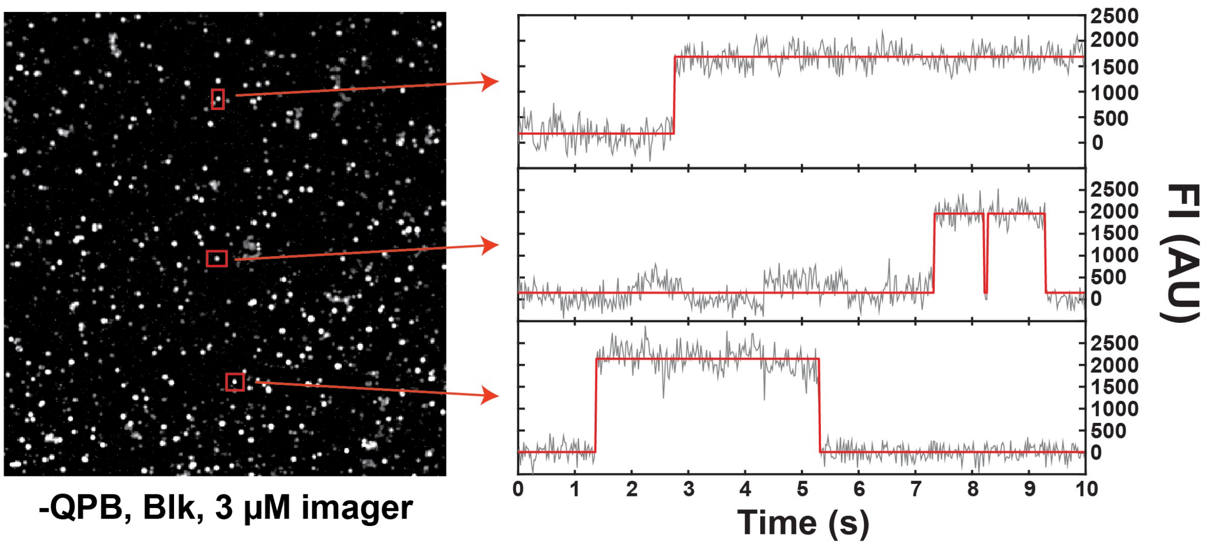

**Supplementary Fig. 2 | Representative traces from non-specifically surface-trapped fluorogenic imagers in the absence of quencher probe blocker (QPB).** A representative blank field of view acquired at 3 µM imager (left) and example fluorescence time traces from selected spots (right). Traces exhibit long-lived, step-like intensity plateaus consistent with surface trapping rather than transient target-binding kinetics (gray, raw; red, HMM idealization).

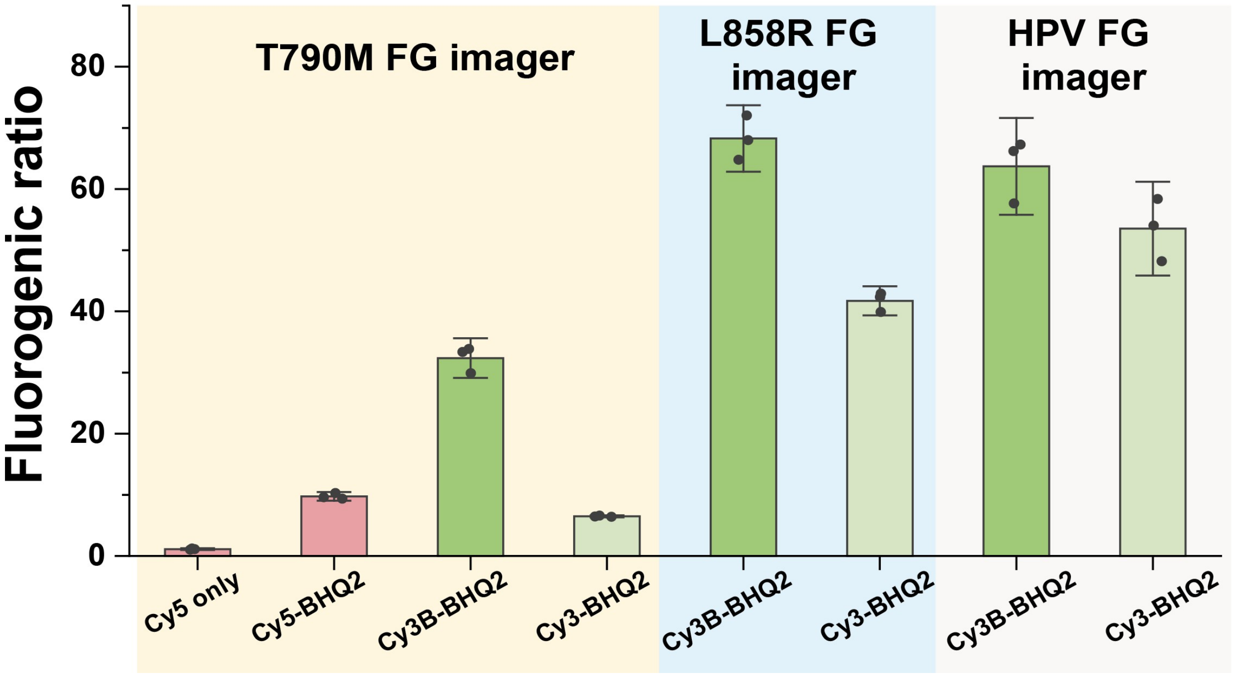

**Supplementary Fig. 3 | Ensemble fluorogenicity of imagers labeled with different fluorophores.** Fluorogenic ratio was calculated as $F = I_{\mathrm{bound}}/I_{\mathrm{unbound}}$. Bars show mean ± s.d. ($n = 3$ independent experiments). Measurements were performed on a SpectraMax ID3 plate reader using clear-bottom 96-well plates. Imagers (1 μM) with or without quencher were mixed with a fully complementary unmodified oligonucleotide (10 μM) in 4× PBS. Fluorescence was measured with 100 ms integration (medium PMT): excitation/emission 530/570 nm for Cy3B- or Cy3-labeled imagers and 630/670 nm for Cy5-labeled imagers. Background signals from 4× PBS were subtracted before calculating fluorogenic ratios.

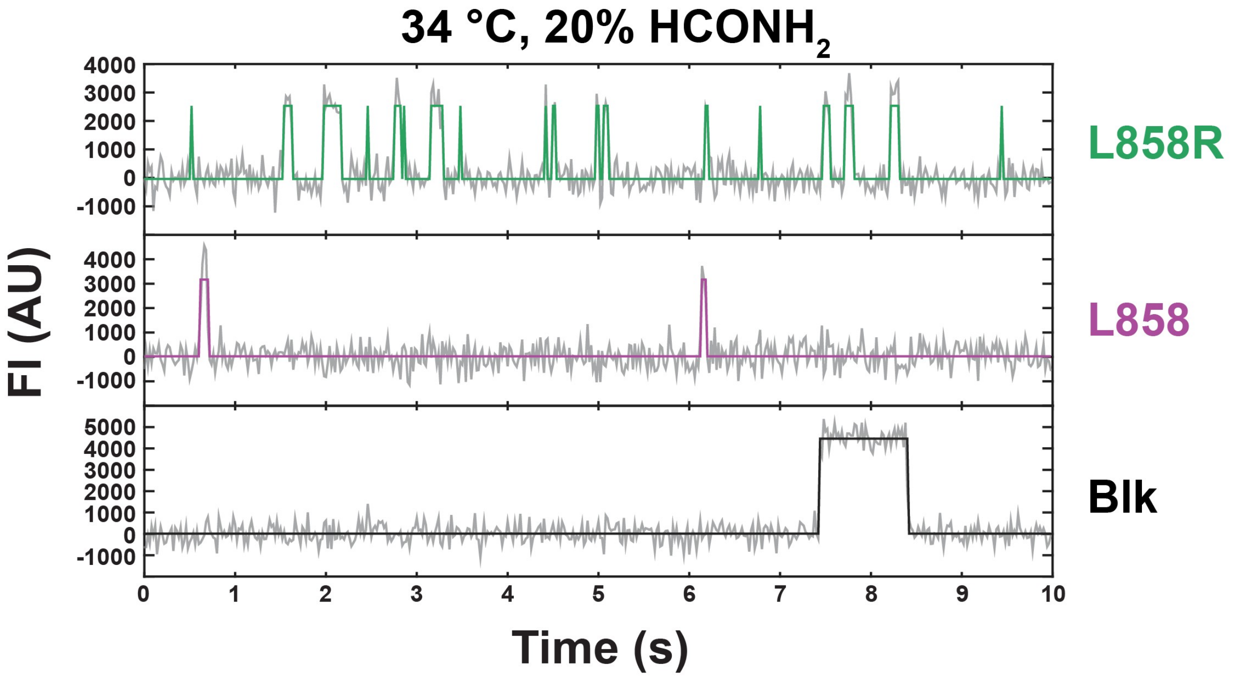

**Supplementary Fig. 4 | Representative kinetic trajectories for *EGFR* L858R, wild-type L858 and blank.** Example single-molecule fluorescence time traces acquired under the optimized conditions used in Extended Data Fig. 4 (34 °C, 20% formamide). Gray, raw trajectories; colored overlays, HMM idealizations.

#### Supplementary Tables

**Supplementary Table 1 | miRNA/DNA sample names and sequences. All sequences are listed 5′-to-3′.**

| Sample Type | Name | Sequence |
|---|---|---|
| Targets | *hsa*-miR-16 | 5´/Phos/rUrArGrCrArGrCrArCrGrUrArArArUrA rUrUrGrGrCrG/3´ |
| | HPV16 26nt_v0 | 5´/ TAGTATAAAAGCAGACATTTTATGCA /3´ |
| | Exon20 T790M 25nt | 5´/ CTCATCATGCAGCTCATGCCCTTCG/3´ |
| | Exon20 WT 25nt | 5´/ CTCATCACGCAGCTCATGCCCTTCG /3´ |
| | *EGFR* L858R MUT 26, Fr | 5´/GTCAAGATCACAGATTTTGGGCGGGC /3´ |
| | *EGFR* L858 WT 26, Fr | 5´/ GTCAAGATCACAGATTTTGGGCTGGC /3´ |
| Capture Probes (CPs) | *hsa*- miR-16 S LNA CP Biotin | 5´/ C+GT+GC+TGC+TA/TEG-Biotin//3´ |
| | HPV16 LNA CP_v1_0 | 5´/Biotin TEG/TGCATAAAATG /3´ |
| | HPV16 LNA CP_v2_0 | 5´/ Biotin TEG/+T+G+C+A+T+A+A+A+A+T+G / 3´ |
| | T790M LNA CP | 5´/ Biotin TEG/CGAAGGGCATG /3´ |
| | T790M LNA CP_v2_0 | 5' / Biotin TEG/CG+A+A+G+G+GC+A+T+G /3' |
| | L858R DNA CP_v1_0 | 5'/ GTGATCTTGAC/3BioTEG/3' |
| | L858R LNA CP_v2_0 | 5'/ +G+T+G+AT+C+T+T+G+A+C/3BioTEG/3' |
| | *hsa*-miR16_NS DNA FP_Cy5_C 10nt | 5'/Cy5/ CGCCAATATT/3' |
| | *hsa*-miR16_NS DNA FP_Cy5_C 10nt | 5'/Cy3/ CGCCAATATT/3' |
| | *hsa*-miR16_FG Imager_v1_0_Cy5 | 5'/5Cy5/ TCGCCCCAATTTAT/3BHQ2/3' |
| | HPV16 FG Imager_v1_0_Cy3B | 5'/ 5Cy3B/TCTGCTCCTACCCTA/3BHQ2 /3' |

| | | |
|---|---|---|
| Fluorescent Probes (FPs) (with Cy5/ Cy3/Cy3B) | HPV16 FG Imager_v1_0_Cy3 | 5'/ 5Cy3/TCTGCTCCTACCCTA/3BHQ2 /3' |
| | T790M_ Cy5 ONLY Fluorogenic Probe V4 | 5'/ /Cy5/AGCTAAATAATGAG /3' |
| | T790M_BHQ2+Cy5 Fluorogenic Probe V4 | 5'/ Cy5/AGCTAAATAATGAG/3BHQ2 /3' |
| | T790M_BHQ2+Cy3B Fluorogenic Probe V4 | 5'/ Cy3B/AGCTAAATAATGAG/3BHQ2/3' |
| | T790M_BHQ2+Cy3 Fluorogenic Probe V4 | 5'/ Cy3/AGCTAAATAATGAG/3BHQ2 /3' |
| | T790M_Cy5 25nt | 5'/ 5Cy5/CTCATCATGCAGCTCATGCCCTTCG /3' |
| | L858R FG Imager_v1_0_Cy3B | 5'/ 5Cy3B/GCTCGCTCTATATCT/3BHQ2 /3' |
| | L858R FG Imager_v1_0_Cy3 | 5'/ 5Cy3/GCTCGCTCTATATCT/3BHQ2 /3' |
| Blockers | *hsa*-miR-16_ S LNA CP Blocker | 5´/ TAGCAGCAC /3' |
| | *hsa*-miR-16_ BHQ 2 blocker | 5´/ AGCGG/3BHQ_2 /3' |
| | HPV16 LNA CPBlocker_v2_0 | 5´/ATTTTATGCA /3' |
| | T790M LNA CP Blocker | 5'/ATGCCCTTCG /3' |
| | L858R LNA CPBlocker_v2_0 | 5'/ GTCAAGATCA /3' |
| Carriers | dT 10 | 5'/TTTTTTTTTT/3' |
| | dT 30 | 5'/TTTTTTTTTTTTTTTTTTTTTTTTTTTTTT /3' |

**Supplementary Table 2 | Capture and imaging conditions used for each FINDER assay.**

| Target (analyte) | Assay type | Capture probe (surface) | Capture temp. | Formamide during capture | Imaging temp. | Formamide during imaging | Imager fluorophore/quencher | Imager conc. |
|---|---|---|---|---|---|---|---|---|
| *hsa*-miR-16 | miRNA | Biotinylated LNA | Room temp. or 37 °C | 0% | 25 °C | 0% | Cy5–BHQ2 | 2 µM |
| HPV16 | Viral DNA | Biotinylated LNA | 37 °C | 0% | 25 °C | 0% | Cy3B (or Cy3)–BHQ2 | 5 µM |
| EGFR T790M vs T790 | SNV | Biotinylated LNA | 37 °C | 0% | 28 °C | 10% | Cy3B–BHQ2 | 5 µM |
| EGFR L858R vs L858 | SNV | Biotinylated LNA | 37 °C | 15% | 34 °C | 20% | Cy3B–BHQ2 | 5 µM |

**Supplementary Table 3 | General parameter sets for trace generation and analysis**

| General Trace Generation Parameters | |
|---|---|
| Spot fixing method | Fluctuation map |
| Spot fixing threshold | 4 |
| Edge Pixel | 20 |
| Start frame | 1 |
| End frame | 750 |
| Percentilecut | 0.8 |
| ROI size (pixels) | 3 |

| General Trace Analysis Parameters | |
|---|---|
| Start frame | 1 |
| End frame | 3000 |
| Exposure time (s) | 0.02 |
| Remove_single_frame_events | True |
| Intensity Threshold | 100 |
| S/N Threshold (Event) | 2 |
| S/N Threshold (Trace) | 3 |
| Minimum $N_{b+d}$ | 10 |
| Minimum $N_{b+d}$ | Inf |
| Minimum $\tau_{on,med}$ | 0.02 |
| Maximum $\tau_{on,med}$ | Inf |
| Minimum $\tau_{off,med}$ | 0.02 |
| Maximum $\tau_{off,med}$ | Inf |
| Maximum individual $\tau_{on}$ | Inf |
| Maximum individual $\tau_{off}$ | Inf |
| Maximum $\tau_{on}$ (C.V.) | Inf |
| Maximum $\tau_{off}$ (C.V.) | Inf |

### Supplementary Notes

#### Supplementary Note 1 | Computational pipeline for fluorogenic probe design for FINDER.

This Supplementary Note describes the computational workflow used to design fluorogenic imager probes for FINDER. The pipeline comprises two MATLAB scripts:

1. **HTfluorogenicScreenV2_3.m**: generation and thermodynamic scoring of a comprehensive library of candidate imager sequences
2. **ThirdRoundOpt_v1_0.m**: score-based clustering, visualization, and selection/optimization of top candidates

##### Part 1. High-throughput screening of candidate sequences (*HTfluorogenicScreenV2_3.m*)

###### 1.1 Initialization and parameter definition

Key design parameters are specified at the outset:

```
Nmm=4;          % Number of mismatches

EndLen5=3;      % Number of bp to leave on 5' imager ends

EndLen3=3;      % Number of bp to leave on 3' imager ends
```

**Parameter rationale:**

**Number of mismatches (Nmm).** We used **Nmm = 4**, which is appropriate for sequences with moderate-to-high GC content and provides useful sequence diversity while preserving discrimination between mutant and wild-type targets. In general, increasing **Nmm** (e.g., 4–5) can increase candidate diversity (often beneficial for lower-GC contexts), whereas decreasing **Nmm**

(e.g., 3) yields fewer, typically more stable binders (often beneficial for higher-GC contexts). Mismatches are introduced to promote discrimination between mutant and wild-type targets.

**Terminal “closing arms” (EndLen5, EndLen3).** We used **EndLen5 = EndLen3 = 3** nucleotides to preserve stable base pairing at both termini of the imager–target duplex. These terminal regions help favor a closed conformation upon binding (supporting fluorophore–quencher separation) and improve structural integrity of the bound duplex.

##### 1.2 Target sequence definition

Mutant and wild-type target sequences are defined (15 nt), and reverse complements are computed to obtain the imager-binding complements:

```
Full='TAGTATAAAAGCAGA';        % Mutant 15 nt full complement

WT='TAGTATAAAAGCAGA';          % WT 15 nt full complement

FullComp=seqrcomplement(Full);

WTFullComp=seqrcomplement(WT);
```

##### 1.3 Reference thermodynamic calculation

Thermodynamics for a validated reference imager (“FP1”) are computed to provide a benchmark for candidate ranking:

```
[FP1dH,FP2dS]=nnHS('CTGCATGA');  % Calculate dH and dS for reference FP1 imager

T=273.15+25;                     % Temperature in Kelvin (25°C)

RTj=2.479;                       % Gas constant in kJ/mol

FP1dG=(FP1dH-FP2dS*T)/RT;        % Free energy of FP1 imager
```

**Purpose.** FP1 provides an experimentally validated kinetic/thermodynamic reference for single-molecule imaging[1]. Candidate ΔG values are compared to FP1 to favor sequences with similar binding/unbinding behavior.

##### 1.4 Combinatorial sequence generation

The candidate library is generated by enumerating mismatch locations within a protected terminal framework and substituting nucleotides at mismatch sites.

**Step 1: Mismatch position combinations.**

```
mmIndList = nchoosek(1:Length-EndLen5-EndLen3,Nmm) + EndLen5;
```

Mismatch indices are drawn only from the **variable internal region**, excluding protected terminal bases defined by *EndLen5* and *EndLen3*. For a 15-nt target with EndLen5 = EndLen3 = 3, the variable region length is 9 nt.

**Step 2: Nucleotide permutations at mismatch sites.**

```
ntV = nchoosek('AGCT',1);

NewList = ntV;

for i = 1:Nmm - 1

   NewList = StrMultiply(NewList,ntV);

end
```

All nucleotide combinations at mismatch sites are generated (nominally 4^Nmm), using a custom string multiplication function, with sequences identical to the original at mismatch positions excluded.

**Step 3: Full library assembly.**

For each mismatch-position combination and each nucleotide permutation, the script:

- constructs a candidate imager sequence,
- enforces uniqueness (no redundant candidates),
- generates a total candidate count of **C(Length − EndLen5 − EndLen3, Nmm) × 3^Nmm**, where **3^Nmm** reflects exclusion of identity substitutions at mismatch sites.

**1.5 Scoring system for candidate evaluation**

Each unique candidate is evaluated using two thermodynamic metrics and a composite score.

**Metric 1: Discrimination energy (ΔΔG).**

```
[dHm,dSm]=nnHSmm(Seq,Full);     % Hybridization with mutant target

[dHwt,dSwt]=nnHSmm(Seq,WT);     % Hybridization with wild-type target

ddG = ((dHm-T*dSm)-(dHwt-T*dSwt))/RTj;
```

ΔΔG reflects preferential binding to mutant vs. wild-type. Candidates with **ddG < 0** (i.e., favoring wild-type) are rejected downstream by assigning a score of zero.

**Metric 2: Reference similarity (ΔG relative to FP1).**

```
dGcompFP = (dHm-T*dSm)/RTj - FP1dG;
```

This term penalizes candidates that deviate strongly from the FP1 thermodynamic baseline, thereby favoring sequences expected to have suitable single-molecule binding lifetimes.

**Composite score.**

Score = (ddG^2 + 1) / (dGcompFP^2 + 1);

This formulation rewards high discrimination (ΔΔG) while penalizing thermodynamic deviation from FP1.

**1.6 Output generation**

Results are saved to a MATLAB *.mat* file containing:

- **AllSeq**: candidate sequence list (cell array)
- **ddG**: discrimination energies (mutant vs. wild-type)
- **dGcompFP**: ΔG deviation from FP1
- **Score**: composite score used for ranking

**Part 2. Candidate selection and optimization (*ThirdRoundOpt_v1_0.m*)**

**2.1 Data loading and initial filtering**

Screening results are loaded and non-discriminating candidates are removed:

load('Screen Result HPV.mat')

Score(ddG<0)=0;

**2.2 Score visualization and cluster identification**

**Score distribution visualization.**

plot(Score,'.')

This plot is used to identify separable groups (“clusters”) of candidates with similar performance.

**Automated clustering boundaries.**

```
C = unique(Score');

dC = diff(C);

UpperBound = C + 0.4*[dC; dC(end)];

BottomBound = C - 0.4*[dC(1); dC];
```

Cluster boundaries are defined using ±40% of the spacing between unique score values to capture natural groupings.

**Cluster selection.**

```
Bnd = Bnd(1:3,:); % Take the first three clusters
```

The top three clusters are carried forward for detailed analysis.

**2.3 Detailed sequence analysis within selected clusters**

For candidates in each selected cluster, the script computes several interpretable sequence features.

**Largest contiguous match length (proxy for continuous complementarity).**

```
props = regionprops(AllSeq{Ind{i}(j)}==FullComp);

largestAreas{i}(j) = max([props.Area]);
```

**Base-pairing propensity factors.**

```
GCpairingFactor{i}(j) = min([sum(AllSeq{Ind{i}(j)}=='G') sum(AllSeq{Ind{i}(j)}=='C')]);

ATpairingFactor{i}(j) = min([sum(AllSeq{Ind{i}(j)}=='T') sum(AllSeq{Ind{i}(j)}=='A')]);
```

Lower GC/AT pairing factors are preferred to reduce the likelihood of stable secondary structure and undesired interactions.

**Base composition.**

```
Gcontent{i}(j) = sum(AllSeq{Ind{i}(j)}=='G');

Acontent{i}(j) = sum(AllSeq{Ind{i}(j)}=='A');
```

**2.4 Sequence logo generation**

Sequence logos summarize positional base preferences within each cluster:

```
seqlogo(AllSeq(Ind{i}))
```

**2.5 Manual optimization process**

Candidates are prioritized by combining cluster score with structural heuristics:

```
cellfun(@min,GCpairingFactor)  % Minimum GC pairing in each cluster

cellfun(@min,ATpairingFactor)  % Minimum AT pairing in each cluster

find((ATpairingFactor{1}==2) & (GCpairingFactor{1}==1))

AllSeq(Ind{1}(ans))  % Display sequences meeting criteria
```

**Selection priorities.**

1. Highest scoring cluster (thermodynamic optimality)
2. Low GC pairing factor (reduced stable structure propensity)
3. Low AT pairing factor (further reduced structure propensity)
4. Balanced nucleotide composition (supporting appropriate duplex stability)

#### Part 3. Final validation protocol

##### 3.1 NUPACK secondary structure analysis

Selected candidates are validated using NUPACK simulations under experimental-like conditions:

- **1 µM** imager + **1 µM** target binding site
- **4× PBS** (reported $[Na^+]$ = **646 mM**)
- **22 °C**

##### 3.2 Validation criteria

**Criterion 1: Minimal secondary structure.**

The target and binding site should remain effectively single-stranded (no competing intramolecular structures that would reduce imager accessibility). Because mismatch-containing imagers bind with reduced energy, competition from target secondary structure is particularly detrimental.

**Criterion 2: Closed-end conformation upon binding.**

Both the 5′ and 3′ ends should be stably base-paired in the bound state to promote the intended closed conformation and maximize fluorophore–quencher separation, thereby optimizing fluorescence enhancement upon binding.

**Criterion 3: Compatibility with the LNA capture probe.**

Heterodimerization between the imager and the LNA capture probe is assessed to ensure no significant off-target interaction that could increase background signal.

##### 3.3 Iterative optimization

If no candidate satisfies all criteria, the pipeline is iterated by:

1. Adjusting terminal arm lengths (**EndLen = 3–5**)

2. Modifying mismatch count **(Nmm = 3–5**)

3. Selecting an alternative imager binding site on the target

4. Re-running screening, clustering, and validation with updated parameters

**Supplementary Note 2 | Determination of required imaging time from a target significance level.**

To determine the required imaging time to meet a given significance level (i.e., separation of the $N_{transition}$ values of the signal and background peaks by a given number of standard deviations), we used the following definition of z-score:

$$z = \frac{\mu_{signal} - \mu_{background}}{\sqrt{\sigma_{signal}{}^2 + \sigma_{background}{}^2}} \tag{1}$$

where $\mu_{signal}$ and $\mu_{background}$ are the mean values for the signal and background distributions, and $\sigma_{signal}$ and $\sigma_{background}$ are the standard deviations of the signal and background distributions, respectively. Assuming Poisson-distributed $N_{transition}$ values, the standard deviation of each distribution is the square root of the mean, giving:

$$z = \frac{\langle N_{transition}\rangle_{signal} - \langle N_{transition}\rangle_{background}}{\sqrt{\langle N_{transition}\rangle_{signal} + \langle N_{transition}\rangle_{background}}} \tag{2}$$

Since $\langle N_{transition}\rangle_{signal}$ and $\langle N_{transition}\rangle_{background}$ are proportional to observation time, they can be related by the proportionality constant $f$ as

$$\langle N_{transition}\rangle_{background} = f\langle N_{transition}\rangle_{signal} \tag{3}$$

Substituting $f\langle N_{transition}\rangle_{signal}$ for $\langle N_{transition}\rangle_{background}$ and rearranging, we obtain the relationship

$$z = \frac{1-f}{\sqrt{1+f}}\sqrt{\langle N_{transition}\rangle_{signal}} \tag{4}$$

To cast this in terms independent of observation time, we can make the following substitution:

$$\langle N_{transition} \rangle_{signal} = k_{bind+dissoc} t \tag{5}$$

where

$$k_{bind+dissoc} = 2 \frac{k'_{bind} k_{dissoc}}{k'_{bind} + k_{dissoc}} \tag{6}$$

is the combined rate constant of binding and dissociation events per unit time[2]. Substitution gives:

$$z = \frac{1-f}{\sqrt{1+f}} \sqrt{k_{bind+dissoc} t} \tag{7}$$

To plot z-score vs. observation time t, we performed Gaussian fitting of the background and signal peaks of the $N_{transition}$ distribution, which yield $\langle N_{transition} \rangle_{background} = 3.38$ and $\langle N_{transition} \rangle_{signal} = 42.22$ at 15 s acquisition time with 0 or 500 fM *hsa*-miR-16, respectively, and hence, we estimate $f = \frac{3.38}{42.22} = 0.080$. To determine $k_{bind+dissoc}$, we performed exponential fitting of the dwell time distributions of the unbound and bound states to estimate $k'_{bind} =$ estimated $k_{bind+dissoc} = 3.22\ s^{-1}$. Plotting the predicted z-score vs. time gives Supplementary Note Fig. 1.

To obtain a significance level of 5σ, the predicted minimum observation time (from rearrangement of equation (7) and substituting $z = 5$) is:

$$t = \frac{25}{k_{bind+dissoc}} \frac{(1+f)}{(1-f)^2} = 9.9\ s \tag{8}$$

This agrees reasonably well with our empirical findings that a 15-s observation time per field of view yields nearly 0 blank counts, and a lower limit of detection than observation times shorter than 10 s. The fact that blank counts are nearly, but not exactly, zero under these theoretically 5σ conditions may be due to non-ideal (i.e., non-Poisson) behavior resulting in a longer tail than expected for the blank $N_{transition}$ distribution.

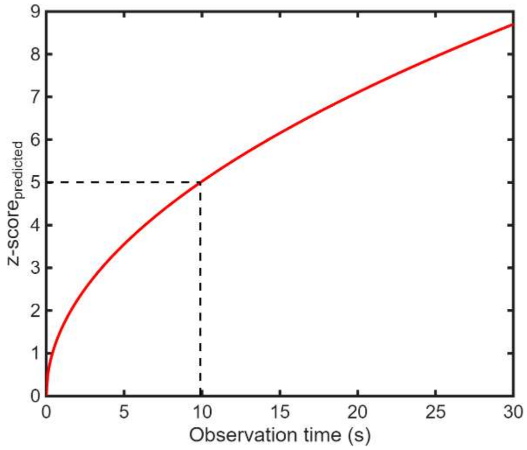

**Supplementary Note Fig. 1| Predicted z-score vs. observation time for the *hsa*-miR-16 assay.** To obtain 5σ separation between the signal and blank $N_{transition}$ distributions, a minimum observation time of approximately 9.9 s is predicted (dashed lines), which is in reasonable agreement with our findings that a 15-s observation time yields nearly perfect separation between signal and background peaks.
